# Heterochromatin organization and liquid-liquid phase separation: it is not about “if” but about “when”

**DOI:** 10.64898/2026.06.03.729812

**Authors:** Hector Romero, Maria Arroyo, Andreas Zhadan, Fernando Muzzopappa, Hui Zhang, Weihua Qin, Marah Mahmoud, Heinrich Leonhardt, Fabian Erdel, M. Cristina Cardoso

**Affiliations:** Cell Biology and Epigenetics, Department of Biology, Technical University of Darmstadt, 64287 Darmstadt, Germany; MCD, Center for Integrative Biology (CBI), University of Toulouse, CNRS, Toulouse, France; Human Biology and Bioimaging, Faculty of Biology, Ludwig Maximilians University Munich, 81377, Munich, Germany

**Keywords:** chromatin modeling, DNA methylation, fluorescence photobleaching (MOCHA-FRAP), heterochromatin, histone methylation, liquid-liquid phase separation, mammalian cells

## Abstract

Heterochromatin is a membraneless compartment within the cell nucleus. In recent years, a controversy arose on whether heterochromatin organization is driven by liquid-liquid phase separation or not. While many heterochromatin proteins were shown to undergo liquid-liquid phase separation *in vitro*, other studies reported that this does not happen in cells. Here, we tested the ability of heterochromatin proteins to generate heterochromatin barrier compartments in cells. We found that, while several proteins (H1.0, H1.4, HP1alpha, HP1beta, Mbd1, Mbd2 and MeCP2) form barrier compartments in mouse and/or human cells this differs between cell types. In addition, not all compartments in the same cell form barriers. We established and experimentally validated a model that predicted the ability to form barrier compartments is dependent on the protein accumulation in heterochromatin followed by the competition between compartments for the nucleoplasm pool of the protein and resulted in larger size for the barrier compartments. These findings resolve the existing controversy and rationalize how in cells heterochromatin compartments form and compete to establish dynamic barriers to the entry and exit of its components.

**Highlights:** Heterochromatin barrier formation differs between proteins, cell lines and heterochromatin compartments within the cell.

Barrier formation depends on heterochromatin anchors, including ligands and other scaffolds.

Barrier compartments are defined by their larger size and higher protein enrichment.

**Graphical Abstract:** 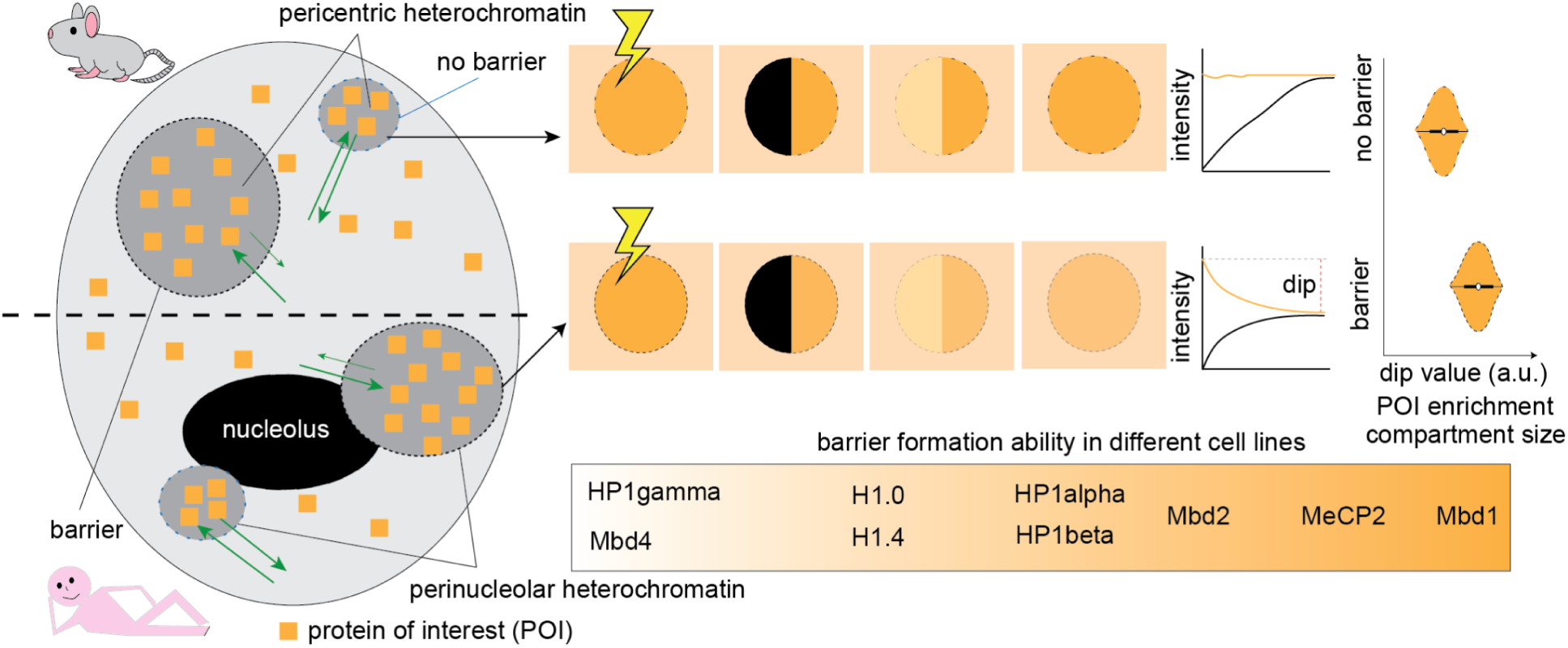

## Introduction

DNA contains all the genetic information that cells require. To be able to fit into the rather small nucleus volume, DNA is compacted into chromatin by wrapping around histone octamers forming nucleosomes. Within the nucleus, the nucleosomes are not distributed evenly, leading to two differentiated phases: euchromatin and heterochromatin. The euchromatin is open enough to allow cellular processes to happen, with special relevance of the gene transcription, while the heterochromatin is highly compacted and correlated to repression. Both compartments are defined by epigenetic markers, located in both the DNA and the histones. With increasing genome sizes in evolution, heterochromatin has greatly expanded, constituting the largest portion of the genome. The major epigenetic markers of heterochromatin are 5-methyl-cytosine (5mC) (Thakur *et al*., 2021) and the trimethylation of residues of the histone 3 (H3K9me3 and H3K27me3) (Guenatri *et al*., 2004). Although many factors that are involved in heterochromatin formation and maintenance have been described, the mechanism of how euchromatin and heterochromatin are separated is not known. In recent years, many of the components of the heterochromatin were found to undergo liquid-liquid phase separation (LLPS) *in vitro*, starting with the nucleosomes (Gibson *et al*., 2019) and the linker histone H1 (Shakya *et al*., 2020). Other components of heterochromatin with LLPS abilities are: HP1alpha (Larson *et al*., 2017; Keenen *et al*., 2021), HP1beta (Qin, Stengl, *et al*., 2021; Qin, Ugur, *et al*., 2021), HP1gamma (Zhang *et al*., 2023; Shen *et al*., 2025), Mbd2 (Zhang *et al*., 2025; Maurici *et al*., 2025) and MeCP2 (Zhang *et al*., 2022; Wang *et al*., 2020; Li *et al*., 2020). However, other studies suggested that HP1alpha LLPS ability was not needed in heterochromatin (Erdel *et al*., 2020), or suggested that the heterochromatin referred to in most of the LLPS studies was limited to certain species including *Mus musculus (Pantier et al., 2024)*. Altogether, this has raised quite some controversy. Importantly, the studies pro and anti-LLPS in heterochromatin are based on different experimental settings and biological materials, making it difficult to have a comprehensive understanding of the disparate outcomes. In this study, we set out to resolve these disparities in the field. Therefore, we utilized MOCHA-FRAP (Muzzopappa *et al*., 2022), a technique that allows the detection of a barrier function in the heterochromatin compartments, and was first used to assess whether HP1alpha undergoes LLPS in mouse cells (Erdel *et al*., 2020). We applied this approach to a panel of heterochromatin-associated proteins: histone H1 (H1.0 and H1.4), HP1s (HP1alpha, HP1beta and HP1gamma), Mbds (Mbd1, Mbd2, Mbd4 and MeCP2) and the synthetic polydactyl zinc finger domain protein MaSat, which binds major satellite DNA repeats (Lindhout *et al*., 2007). Moreover, the experiments were performed across multiple cell lines from different species including the ones used in previous studies and expanding from them. In addition, we quantified and manipulated the binding sites for these factors in cells. Our results show that most of these proteins are capable of forming barriers within heterochromatin in cells under certain conditions including their cognate binding sites. Additionally, our modeling indicates that there is a competition between compartments enriched in molecular anchors in the cell nucleus for the free proteins available in the nucleoplasmic pool to establish barrier compartments.

## Methods

### Cells

All cells used were tested for mycoplasma and deemed free of contamination. Cell line characteristics and description are depicted in Table S1.

C2C12 myoblasts were cultured in high-glucose Dulbecco′s Modified Eagle′s Medium (DMEM) (Sigma-Aldrich Chemie GmbH, D6429) supplemented with 20% fetal calf serum (FCS), 1× L-glutamine (Sigma-Aldrich Chemie GmbH, G7513), and 1 µM gentamicin (Sigma-Aldrich Chemie GmbH, G1397).

The embryonic fibroblast NIH-3T3, MEF W8 (wild type), and MEF D5 (Suv39h1^-/-^ Suv39h2^-/-^), as well as L929 connective tissue fibroblast, MTF tail fibroblast, NDF neonatal human dermal fibroblast and hTERT immortalized human RPE-1 retinal pigment epithelial cells, were cultured in DMEM high glucose supplemented with 10% FCS, 1× L-glutamine and 1 µM gentamicin.

The embryonic fibroblast MEF-P (P53^-/-^), MEF-PM (P53^-/-^ Dnmt1^-/-^) were cultured in DMEM high glucose supplemented with 15% FCS, 1× L-glutamine, and 1 µM gentamicin.

For subculturing of myoblasts and fibroblasts, the media was aspirated, cells were briefly washed using 0.02% EDTA in PBS and then incubated in trypsin-EDTA (Capricorn Scientific, TRY-3B) for 5 min at 37 °C and 5% CO_2_. The reaction was stopped by adding at least twice volume of growth media, and then added to a new plate containing fresh growth media.

The J1-derived neural stem cells (NSC) were cultured in plates, coverslips or slide chambers coated with D-lysine and laminin. The coating was settled by incubation in 10 µg/ml poly-D-Lysine (Sigma Aldrich, P7405) in H_2_O for 4 h followed by drying at room temperature for 20 min and an overnight incubation at 37 °C with 5 µg/ml laminin (Sigma Aldrich, L2020) in ice-cold DMEM:F12 medium (Sigma Aldrich, 56498C). NSC were cultured in media composed of Euromed-N (Biozol Diagnostica Vertrieb, ECL-ECM0883L), 1× N-2 supplement (ThermoFischer Scientific, 17502048) 1× L-glutamine, 1× penicillin/streptomycin (Sigma Aldrich, P0781), 20 ng/ml murine fibroblast growth factor-2 (Peprotech, PPT-450-33-500) and 20 ng/ml murine epidermal growth factor (Peprotech, 315-09-500UG). For subculturing, the media was aspirated, and the cells were briefly washed with PBS before incubation with Accutase (Sigma Aldrich, A6964) for 3 min. The reaction was inactivated by addition of 2× volumes of growth media and transferred to a new coated vessel containing fresh media.

### Transfection

For transfection, cells were detached from the vessel as described above and centrifuged at 1400 rpm for 5 min. Media was aspirated and cells were resuspended in AMAXA M1 buffer (5 mM KCl, 15 mM MgCl_2_·6H_2_O, 120 mM Na_2_HPO_4_/NaH_2_PO_4_, 50 mM Mannitol) containing 2-10 µg of plasmid and electroporated using AMAXA nucleofection system (Lonza), using the AMAXA program described in Table S1, and then added to self-made glass-bottom slide chambers containing high precision glass (for MOCHA-FRAP) or plates containing glass coverslips (for immunostaining).

### Plasmids

All plasmid characteristics, as well as the first characterization, are described in Table S2. All plasmid sequences were checked with full plasmid sequencing.

### MOCHA-FRAP

Cells were prepared as described before. In a Leica SP5-II confocal microscope (Table S3), with 100X immersion oil objective, and an additional 40X zoom, half of the heterochromatin compartment was set to bleach using a point ROI function, with maximum intensity of the correspondent laser for 100 ms, so approximately half of it was bleached. Frame rate of acquisition was 700 Hz, leading to a final exposure time of approximately 0.1 s. The pre-bleach was documented by taking a frame before bleaching, followed by 100 to 300 frames of the recovery (10 to 30 s).

The analysis of the MOCHA-FRAP was done as described before (Erdel *et al*., 2020; Muzzopappa *et al*., 2022). In this work, a self-written macro (full-Barista), available in TUdatalib, was used to segment the heterochromatin compartment, the bleach and non-bleach halves of it as well as a reference unbleached heterochromatin compartment, as well as to obtain the intensities from these compartments and the enrichment on the heterochromatin of interest, defined as the mean intensity on the compartment divided by the intensity out of the regions segmented before.

The intensities were normalized following the steps of (Erdel *et al*., 2020; Muzzopappa *et al*., 2022) in a MATLAB GUI (MOCHA_FRAP_GUI_2.m) also available in TUdatalib. The interface allows the visualization of the individual compartments and its classification into barrier (LLPS) and non-barrier, as well as calculating the dips and performing the average curves for each condition. The mean and 95% confidence interval for the average curves were obtained from the extracted csv of the average curves, in the minimum point of the fit.

### Immunofluorescence

For the immunofluorescence staining of H3K9me3, cells were seeded on gelatin-coated coverslips. Cells were washed once with 1x PBS and subsequently fixed in a 3.7% formaldehyde solution for 10 minutes. After fixation, the cells were permeabilized with 1x PBS supplemented with 0.5% Triton X-100 for 20 minutes at room temperature. After washing three times with PBS-T (1x PBS + 0.01% Tween-20), cells were blocked with 1% BSA in PBS-T for 1 hour. Primary and secondary antibodies were diluted in the blocking solution and incubated for 60 minutes. Between and after the two incubation steps, coverslips were washed three times with PBS-T for 5 minutes. DNA was counterstained with DAPI (10 mg/ml, (4’,6-diamidino-2-phenylindole, Sigma-Aldrich Chemie GmbH, Steinheim, Germany, Cat.No.: D9542) for 10 minutes. Coverslips were mounted on glass slides with Mowiol® 4-88 (Sigma-Aldrich Chemie GmbH, Steinheim, Germany, Cat.No.: 81381). All antibody characteristics are summarized in Table S4.

To detect 5mC *in situ*, we followed the instructions of a previously published protocol (Arroyo *et al*., 2023). In brief, the seeded cells were washed once with 1x PBS and fixed in 3.7% formaldehyde solution for 10 minutes. Following the fixation, permeabilization was done with 0.5% Triton X-100 for 20 minutes at room temperature. After washing the permeabilized cells three times with 1x PBS, they were incubated in ice-cold 88% methanol for 5 minutes. The cells were rehydrated for 15 minutes with 1x PBS at room temperature. The cells were treated with 100µg/ml RNaseA in 1x PBS for 1h at 37°C. Subsequently, the cells were washed three times with PBS-T and blocked in 1% BSA in PBS-T for 1h. The cells were incubated in primary antibody solution containing 1% BSA, 1x DNaseI reaction buffer (10 mM Tris-HCl pH 7.5, 2.5 mM MgCl_2_, 0.5 mM CaCl_2_), 0.5 U of DNaseI (Sigma-Aldrich Chemie GmbH, Steinheim, Germany, Cat.No.: D5025) and anti 5mC-antibody (1:250) for 70 minutes at 37°C followed by three wash steps with PBS-TE (PBS-T with 100 mM EDTA). Secondary antibodies were diluted in 1% BSA/PBS and cells were incubated for 1h at room temperature. DNA was counterstained with DAPI as described in the previous paragraph.

#### High-content microscopy

High-content microscopy and analysis were done in the Operetta high-content screening system (Perkin Elmer, UK, Table S3) in wide-field mode, equipped with a Xenon fiber optic light source and a 20x/0.45 NA long working distance or a ×40/0.95 NA objective. For excitation and emission, following filter combinations were used, 360-400 nm and 410-480 nm for DAPI, 460-490 nm and 500–550 nm for GFP as well as 560–580 nm and 590–640 nm for Cy3. Fluorescence intensity levels were quantified with the Harmony software (Version 3.5.1, PerkinElmer, UK). For the analysis of cells transfected with GFP fusion encoding plasmids and stained against H3K9me3 or 5mC and counterstained with DAPI, cell nuclei were first identified according to their DAPI fluorescence and evaluated for morphological properties like roundness and size. In the nuclei fitting the criteria, total intensity and standard deviation of the intensity was calculated for DAPI, GFP (when transfected), and Cy3 labelled antibodies. The intensity of Cy3 was then normalized by dividing them to the intensity of DAPI in each nucleus to compensate for potential cell cycle-dependent fluctuations.

### Molecular dynamics simulation

We used polymer simulations to model generic cases of multivalent proteins binding to clustered heterochromatic regions. To model clustered heterochromatic regions, we simulated six polymers attached to opposite faces of the simulation box. Each polymer represents a chromosomal region containing the heterochromatin, composed of a spacing region of 20 neutral beads followed by the heterochromatic region composed of 50 alternating neutral and heterochromatic beads. Each bead represents 10 kb of chromatin and has a dimensionless radius of σ. Depending on the simulation, 500, 1000 or 5000 proteins of radius σ were added. The chromatin and protein motions were calculated using LAMMPS (Thompson *et al*., 2022) in a simulation box with reflective boundary conditions and an edge length of 100 σ. The chromatin beads were connected by a finitely extensible non-linear elastic (FENE) bonds, with an energy calculated as:

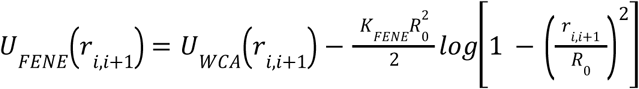

where r(i,i+1) was the separation of the neighboring beads. We used a bond energy K_FENE_ of 30 kBT and a maximum bond extension bond, R_0_, of 1.6 σ. The first term was the Weeks-Chandler-Andersen (WCA) potential:

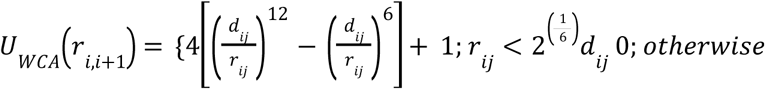

where d_ij_ is the average diameter of beads i and j.

The rigidity of the chromatin polymer was defined by a Kratky-Porod potential:

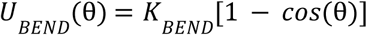

where θ is the angle between three adjacent beads and K_BEND_ is the bending energy. The latter relates to the persistence length of the chromatin fiber l_p_ = K_BEND_/k_B_T. It was defined such that l_p_ equals 1.5-times the diameter of a chromatin bead.

The interaction between the beads was calculated using a truncated Lennard-Jones potential:

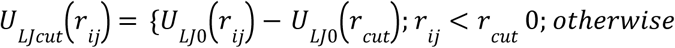

with:

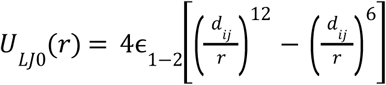

Where r_cut_ is the cut-off distance (r_cut_=2.5σ) and ε_1-2_ is the energy scale of interaction between bead_1_ and bead_2_.

First, the polymers were equilibrated using soft potentials. Then, a Lennard-Jones potential between heterochromatic beads was applied with an energy ε_1-2_ of 1.5 k_B_T, and the collapse and clustering of the heterochromatin was recorded for 0.5 million simulation steps until the radius of gyration of the chromatin was stabilized. Finally, interactions between protein beads and heterochromatic beads, as well as interactions between protein beads and protein beads interactions were simulated by applying an additional Lennard-Jones potential with a ε_1-2_ of 2.25 k_B_T, and the evolution of the system was recorded every 5000 steps for 22 million simulation steps.

To analyze the simulations, we used a custom python script to extract all beads positions from each simulation. For each heterochromatic region, the radius of gyration was calculated as the average distance of each heterochromatic bead from the geometrical center of the region. Then, a distance cutoff of 1.25σ was used to identify proteins directly bound to the heterochromatic beads. The second layer of proteins bound to the direct binders was identified as indirect binders. Heterochromatic regions with more than 80 indirect binders contain more than double of indirect than direct binders. Since this indicates a multivalent-protein mediated mechanism, characteristic of phase-separated condensates, we designated them as compartments with barrier. The percentage of proteins recruited to the heterochromatic region through multivalent interactions was calculated as the ratio of indirect binders to all proteins in the region of interest. Finally, virtual microscopy images were generated from each simulation step by projecting the XY-coordinates of all beads onto a two-dimensional matrix of all XY beads coordinates, and assigning an arbitrary fluorescence intensity value to each bead. The resulting distribution was convolved with a Gaussian point-spread function to generate virtual fluorescence micrographs. The enrichment of proteins at each heterochromatic region was calculated as the median intensity in the structure divided by the median intensity of the nucleoplasm.

### Protein interaction network analysis

Protein interaction networks for HP1 proteins were generated using the STRING database (Szklarczyk *et al*., 2023). For HP1α, the network type was set to “full STRING network,” with network edges interpreted based on confidence. The minimum required interaction score was set to the highest confidence level (0.9), and the maximum number of interactors displayed in both the first and second shells was limited to 20. For HP1β, the network type was set to “physical subnetwork,” with network edges interpreted based on confidence. The minimum required interaction score was also set to the highest confidence level (0.9), while the maximum number of interactors displayed in the first and second shells was limited to 20 and 50, respectively. All remaining parameters were maintained at their default settings. In the resulting interaction maps, colored nodes represent the query proteins and proteins closely associated with them, whereas white nodes indicate secondary interactors or proteins automatically added by STRING. Blue edges between nodes indicate protein-protein interactions.

### Analysis and statistics

For high-throughput data, the outliers were detected and removed using the interquartile range (IQR) in a MATLAB script (Outliers_IQR.m) available in TUdatalib. In this method, the IQR is calculated as the result of Q3 - Q1, and an outlier point is defined as those that fall below Q1 - 1.5 IQR or above Q3 + 1.5 IQR. This was done for each individual dataset.

The datasets were then imported to a MATLAB GUI, where they were assigned to a condition to generate a violin plot and perform statistics between conditions. In the violin plot, the median (white dot), IQR (dark-gray box) and standard deviation (dark-gray line) of the combined datasets are always shown. The mean of each dataset is added as a diamond located over the violin. In the case of the dip, the barrier and non-barrier were separated as different datasets to obtain the mean of each. The statistical comparison of conditions is done based on the means of the replicates using the Welch t test and Hedges g to estimate the value of the differences.

## Results and discussion

### Mouse heterochromatin showed a barrier behavior especially for methyl-CpG binding proteins

For a comprehensive analysis of the ability of heterochromatin proteins to create barriers that dynamically separate the compartment from the rest of the nucleus, we used the MOCHA-FRAP technique, a live-cell approach combining fluorescence recovery after photobleaching (FRAP) and fluorescence loss in photobleaching (FLIP) measurements on the same heterochromatin compartment. This technique has been shown to be a functional tool for assessing barrier formation in liquid-liquid phase separation (Joyot *et al*., 2026). The analysis of the half-bleached compartments have three possible outcomes depending on the strength of the barrier formed (Figure 1A) (Erdel *et al*., 2020): i) when no barrier is formed, the protein can diffuse freely and both the bleach and the non-bleach half can recover at the same pace, therefore producing a very small to no dip in the final graph; ii) when there is a semi-permeable barrier, there will be a first wave of movement from the non-bleached half to the bleached half that is visible as a dip, but later both sides will be able to recover both sides from the exterior; iii) in the presence of an almost impermeable barrier, most of the recovery on the bleached half will come from the non-bleached half, and almost no recovery will come from outside, producing the dip in the form of a plateau.

**Figure 1.**
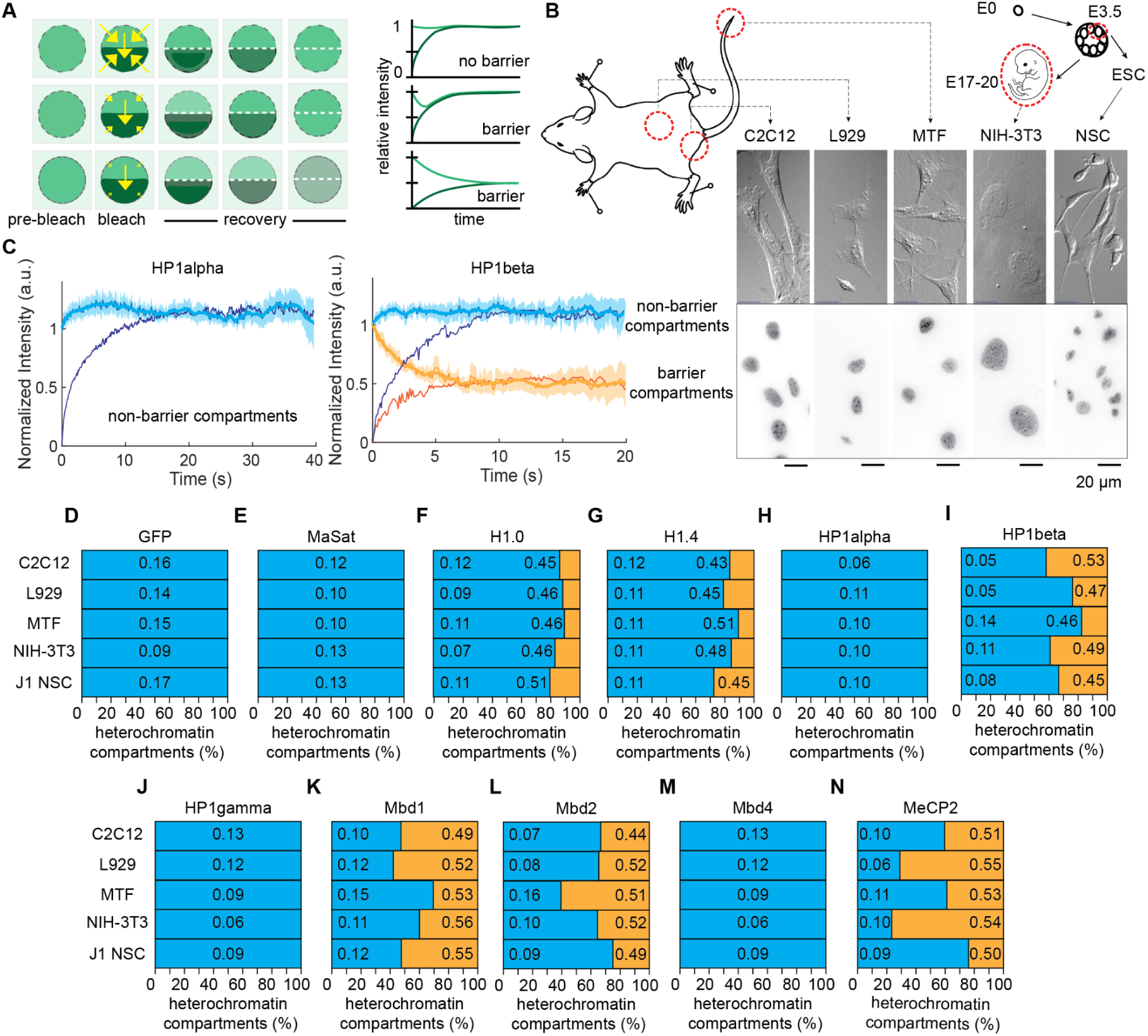
Barrier-forming heterochromatin compartments are mostly driven by Mbds in mouse cells. A. Theoretical background of the MOCHA-FRAP technique. When a POI is enriched in the heterochromatin and half of the compartment is bleached, there are three possible outcomes: when the protein can freely diffuse through the heterochromatin compartment (upper panel), the recovery will take place from any direction, leading to a situation with very small dip indicating no LLPS; if the compartment is partially (middle panel) or almost totally impermeable (down panel) due to the LLPS, the recovery of the bleached half will come mainly from the non-bleached half, leading to a dip. Depending on the permeability of the POI to the compartment, the intensity can be fully recovered in both sites, or just be equivalent. Yellow arrows indicate movement of molecules (size accounts for number of molecules). Modified from Erdel et al. (Erdel *et al*., 2020). B. Mouse cell lines used for the study. C2C12 myoblast are adult stem cells from muscle tissue; L929 are fibroblasts derived from connective tissue; MTF are fibroblast cells derived from the tail of adult mice; NIH-3T3 are fibroblasts from embryonic origin; and J1 NSC are neural stem cells derived from embryonic stem cells by *in vitro* differentiation. The upper panel for each cell line represents a different interference contrast (DIC) image and the lower panel the DNA staining (DAPI) of the same field. C. Exemplary curves for the two types of results obtained in the MOCHA-FRAP screening. Cells were transfected with a plasmid containing the coding region of the gene of interest fused to the GFP gene. Live cells were then visualized under the microscope and 2-5 half heterochromatin bleach and recovery movies were recorded per cell. The recovery was then analyzed in the bleach and non-bleach half to obtain the MOCHA-curves and were analyzed individually based on the shapes described in panel A. In some cases, as HP1alpha (left panel), all compartments were homogenous and showed no barrier, while other proteins as HP1beta (right panel), showed a dual behavior with some compartments in which the non-bleached half had a dip indicating a barrier (orange lines), while others did not (blue lines). Shades represent the 95% confidence interval of the non-bleach curves. D-N. Classification of the heterochromatin compartments for different proteins of interest in no-barrier (blue) or barrier (orange) compartments as a percentage. Proteins of interest are indicated above the plots. In those that have non-barrier and barrier compartments, left hand side numbers correspond to the mean dip of the individual non-barrier compartments (blue) and the right hand side numbers to the mean dip of the individual barrier compartments (orange).

We also looked for cell lines with different origins and characteristics to address the differences in the epigenetic landscape of the heterochromatin. We selected five different mouse cell lines. Three fibroblasts with different origins, connective tissue (L929), dermal epithelium of adult mouse tail (MTF) and embryonic (NIH-3T3), as well as two adult stem cells, from the skeletal muscle tissue (C2C12 myoblasts) and neural stem cells (NSC) derived from embryonic stem cells by *in vitro* differentiation (Figure 1B). In all cells, we studied the behavior of 10 proteins related to heterochromatin. These include histone H1, concretely the isoforms H1.0 and H1.4, both known to accumulate in heterochromatin (Schmidt *et al*., 2024), the three isoforms of the heterochromatin protein 1 (HP1) alpha, beta and gamma that recognize histone H3 K9 trimethylation, the methyl-CpG binding proteins (Mbd) Mbd1, Mbd2, Mbd4 and MeCP2, and the artificially generated zinc finger domain protein MaSat that binds major satellite DNA sequences. We used GFP as a control for a protein not enriched at heterochromatin. For the screening, we transfected a plasmid containing the coding region of the gene of interest fused to GFP (in some cases to RFP), and took images of 40-100 heterochromatin compartments from cells with different protein levels in different biological replicates. We compared that the output GFP fluorescence between the different plasmids used was similar in the positive cells (Figure S1). In the analysis, we noted that those compartments located in cells with very low protein levels had to be discarded as the changes in fluorescence caused by bleaching during imaging masked the dip explained above. This, however, happened for all proteins studied.

The analysis of the screening revealed two groups of proteins. In one group, as shown for HP1alpha (Figure 1C, left panel), all compartments did not show a dip and were considered non-barrier. In a second group, represented by HP1beta (Figure 1C, right panel), some compartments were not having a barrier, while other compartments showed a clear dip. Therefore, we classified them and calculated the dip for each compartment independently (Figure 1D-N). By doing this, we showed that indeed the values of the dip were similar between all proteins, being the non-barrier average values between 0.05 and 0.16 and the barrier ones between 0.43 and 0.55 (Table S5). The most differences became clear from the percentages of compartments that showed a dip: i) GFP (Figure 1D), MaSat (Figure 1E), HP1alpha (Figure 1H), HP1gamma (Figure 1J) and Mbd4 (Figure 1M) showed only non-barrier compartments; ii) H1.0 (Figure 1F) and H1.4 (Figure 1G) showed consistent low percentages (10-20%) of barrier compartments throughout the different cell lines; iii) HP1beta (Figure 1I) showed low to medium percentages (20-50%) of barrier compartments); iv) Mbd1 (Figure 1K), Mbd2 (Figure 1L) and MeCP2 (Figure 1N) showed highest percentages of barrier compartments, with more than 50% of them in some cases. They also showed the highest variability among cell lines, especially Mbd2 and MeCP2. The results of the screening match and extend the already reported behavior of HP1alpha (Erdel *et al*., 2020) as it does not form barrier compartments in any of the cell line studied.The positive barrier formation reported in the studies with stronger *in cellulo* evidence for histone H1 (Shakya *et al*., 2020), MeCP2 (Zhang *et al*., 2022) and Mbd2 (Zhang *et al*., 2025) are also validated and expanded to cells of different origins. In contrast, Mbd1 and Mbd4 show different behaviors in barrier formation although both are not able to undergo LLPS *in vitro* (Zhang *et al*., 2025), suggesting that Mbd1 requires additional factors not provided in the *in vitro* assays to generate the barrier. Even in the cases with stronger barrier formation, there is a fraction of compartments where the barrier was not formed, and these percentages were largely conserved among the different cell lines studied.

### Barrier compartments are rare but exist in human cells

Next, we tested if the barrier formation was, as it was mentioned before (Pantier *et al*., 2024), an unique feature of mouse cells. Therefore, we used MOCHA-FRAP to screen the same proteins in two (diploid) human cell lines (Figure 2A): immortalized retinal pigment epithelium (hTERT-RPE1) and neonatal dermal fibroblasts (NDF). These cell lines are diploid, stable and tend to form perinucleolar heterochromatin compartments that are comparable to those in the mouse and, therefore, amenable for the MOCHA-FRAP technique. In these human cell lines, the dip for the non-barrier compartments were slightly smaller than in the mouse (0.02-0.13), while the barrier compartments showed similar values (0.42-0.57) (Figure S3, Figure 2B-L, Table S6). In both cell lines, MaSat could not enrich in heterochromatin (from ∼3-fold in mouse to ∼1.2-fold in human cells) (Table S7), therefore it was cotransfected with HP1alpha to visualize the compartments. In the case of MeCP2, HP1alpha was co-expressed in hTERT-RPE1 as MeCP2 accumulation in heterochromatin was rare, but this was not the case in NDF cells, even though the enrichment dropped from values 7-10 in mouse cells to 2.0 and 2.6 in hTERT-RPE1 and NDF, respectively (Table S7). The results of the screening showed that: i) GFP (Figure 2B), MaSat (Figure 2C), HP1gamma (Figure 2H) and Mbd4 (Figure 2K) showed only non-barrier compartments as was the case in mouse cells; ii) H1.0 (Figure 2D), H1.4 (Figure 2E) and Mbd1 (Figure 2I) had comparable percentages of barrier compartments to the ones exhibited in mouse cells; iii) HP1alpha (Figure 2F) showed low to middle percentages of barrier compartments; iv) HP1beta (Figure 2G), Mbd2 (Figure J) and MeCP2 (Figure 2L) lost partially or totally the ability to form barrier compartments.

**Figure 2.**
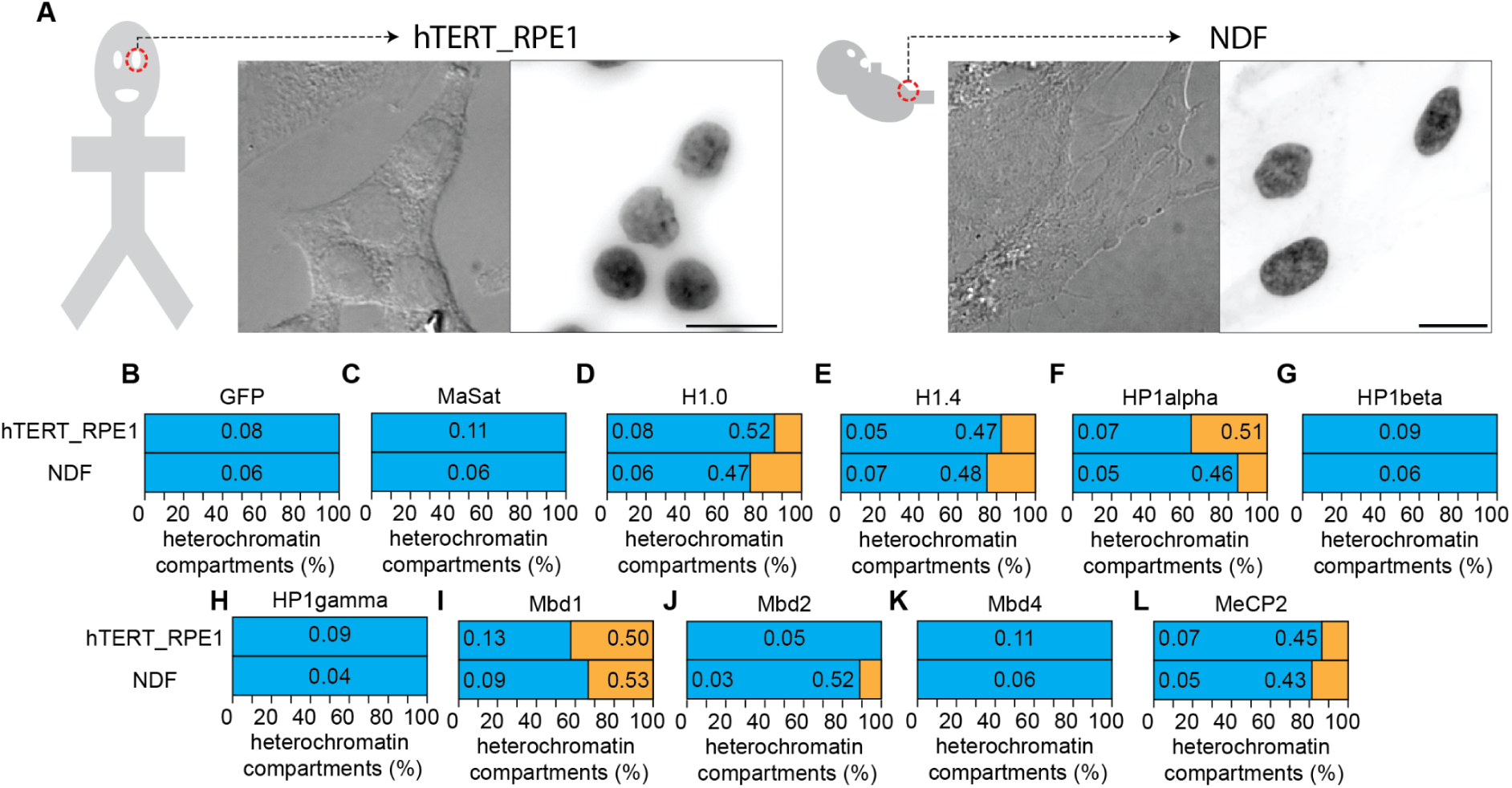
Barrier compartments are rare and limited to few proteins in human cells. A. Human cell lines used in the study and their origin. hTERT-RPE1 are cells derived from retinal epithelium immortalized by transfection of the telomerase reverse transcriptase (hTERT) gene and NDF are immortal cells derived from neonatal dermal fibroblasts. The left panel shows a differential interference contrast (DIC) image, and the right panel DNA staining (DAPI). B-L. MOCHA-FRAP screening performed in the human cell lines for the protein of interest as indicated above the plots. Blue represents non-barrier compartments and orange barrier compartments. When both are present, left hand side numbers represent the average dip from the individual non-barrier compartments and right hand side numbers the average dip from the individual barrier compartments.

This second screening in human cells, thus, revealed that MeCP2 barrier formation was residual compared to the mouse cell lines (Figure 1N), but than rather than being a unique feature of *Mus musculus* (Pantier *et al*., 2024), there must be underlying conditions in the mouse cells used that enhance the barrier formation ability of MeCP2. In fact, likely for this reason, mouse models are often used to characterize the functions of this protein (Ito-Ishida *et al*., 2020; Heckman *et al*., 2014; Singleton *et al*., 2011; Romero *et al*., 2025; Zhang *et al*., 2022; Guy *et al*., 2001; Tillotson *et al*., 2017). As Mbd2 followed the same pattern, this could be related to the 5mC distribution on these heterochromatin compartments. Nonetheless, Mbd1 barrier formation was mostly unaltered. Hence, we used the short isoform of Mbd1 (Mbd1ΔCxxC), which lacks the CxxC3, responsible for unmethylated CpG binding (Figure S4A) (Jørgensen *et al*., 2004) in the different mouse and human cell lines. Mbd1ΔCxxC behaved like Mbd1 in all mouse cell lines (Figure S4B), but failed to produce barrier compartments in both hTERT-RPE1 and NDF cells (Figure S4C), most strongly in hTERT-RPE1 (Figure S4D). This suggests that it is indeed a 5mC effect, probably due to the fact that the CpG islands in the perinucleolar compartments are not methylated enough.

Also interesting is the change of behavior of HP1alpha and HP1beta between human and mouse cell lines. These differences could be related to the interactomes of HP1alpha and beta in mouse and human cells (Figure S5). Specifically, according to the STRING database (Szklarczyk *et al*., 2023), Shugoshin-1 (SGO1) and HP1alpha (Cbx5) are enriched in the human HP1beta interaction network (Figure S5B), but not similarly enriched in mouse cells (Figure S5A). SGO1 interacts with the HP1s through a basic region (Larson *et al*., 2017), which could block the acidic region of HP1beta required for liquid-liquid phase separation (Qin, Stengl, *et al*., 2021) and potentially limit its barrier formation ability. In addition, HP1alpha enrichment in the human heterochromatin is twice as high as in the mouse heterochromatin (Table S7), which could act both as an activation for its barrier formation and limit the H3K9me3 ligand for HP1beta, another factor required for LLPS (Qin, Stengl, *et al*., 2021) and therefore limiting its barrier formation ability. HP1alpha mouse interactome (Figure S5C) includes Lamin B receptor (Lbr), while human HP1alpha does not (Figure S5D) and included instead HP1beta (Cbx1). Although both Lbr and HP1beta have been reported to enhance HP1alpha LLPS ability (Larson *et al*., 2017; Phan *et al*., 2024; Keenen *et al*., 2021), there might be differences in the extent of the inhibition that allowed HP1alpha to form barrier in human cells but not in mouse cells.

### HP1beta and MeCP2 abilities for heterochromatin barrier formation are regulated by their ligands

We wondered if the differences in the barrier versus non-barrier compartments between cell lines had a relation to the way the proteins bind the heterochromatin. As depicted in Figure 3A, there are five types of binding in the proteins studied: i) GFP is not interacting at all, therefore is not enriched in heterochromatin; ii) MaSat binds to the major satellite DNA repeats (Lindhout *et al*., 2007), which in mouse are mostly located in the pericentric heterochromatin; iii) histone H1 bind to the entry/exit points of the DNA wrapped around nucleosomes (Cutter and Hayes, 2015), being H1.0 and H1.4 naturally enriched in heterochromatin (Th’ng *et al*., 2005); iv) the HP1s bind to the trimethylated lysine 9 of the histone H3 (H3K9me3) (Machida *et al*., 2018); v) Mbds bind 5-methyl cytosine (5mC) with preference for methyl-CpG sequences (Liu *et al*., 2018), but they have additional DNA binding motifs that differ among the Mbd proteins (Zhang *et al*., 2017).

**Figure 3.**
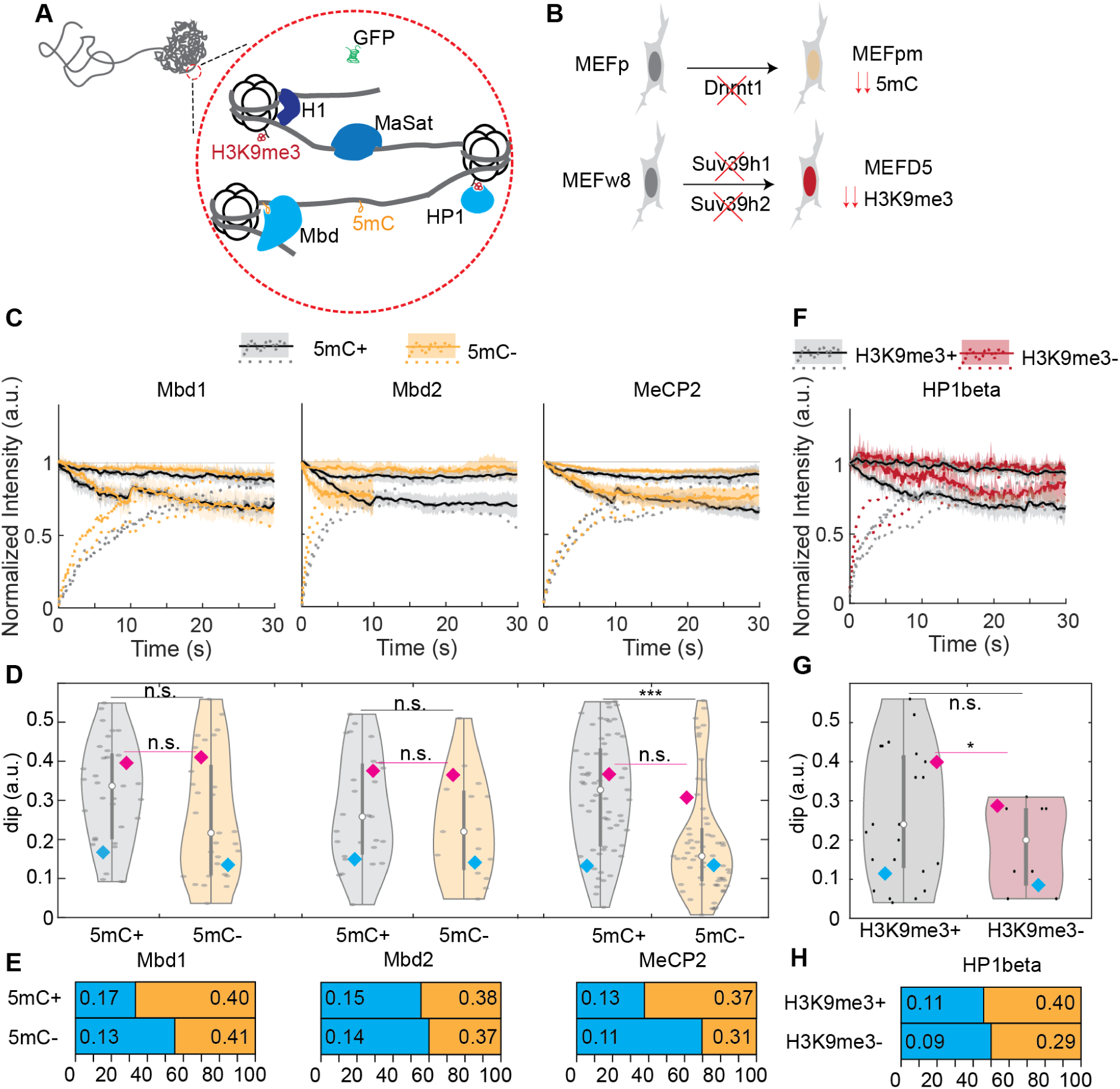
HP1beta and MeCP2 heterochromatin barrier formation is regulated by their ligands H3K9me3 and 5mC. A. Summary of the different binding factors of the proteins studied. GFP does not have any interaction and diffuses freely within the cell; MaSat binds to major satellite repeats, which are mostly located in the pericentric heterochromatin in mouse; H1s bind to the nucleosome DNA entry/exit site, interacting with both linker DNA and the core histones; the Mbds bind to methylated CpG due to its main ligand, 5 methyl-cytosine, which are often located in or close to the triad of the nucleosome; HP1s bind to its main ligand, the trimethylated lysine 9 of the histone H3 tail (H3K9me3). B. Cell lines used to determine the effect of the ligands in the barrier formation. MEFp are embryonic fibroblasts with the genetic background *p53^-/-^*, which is necessary when knocking out the *Dnmt1* gene in MEFpm. As Dnmt1 maintains the 5mC during replication, the knock-out shows very reduced levels of 5mC. MEF w8 is a wild type strain of mouse embryonic fibroblasts used for generating the double knock-out *Suv39h1* and *Suv39h2* named MEF D5 These proteins are responsible for the methylation of the lysine 9 of H3, therefore the levels of H3K9me3 are very low in these cells. C. Average MOCHA-FRAP curves from Mbd1 (left panel), Mbd2 (middle panel) and MeCP2 (right panel) on the 5mC proficient MEFp (black and gray) and 5mC deficient MEFpm (yellow). Each compartment was analyzed alone to determine the belonging to barrier or not barrier and then averaged. Lines represent the average of all compartments within the same category and shades the 95% confidence interval. D. Violin plot of the dips calculated for Mbd1 (left), Mbd2 (middle) and MeCP2 (right) in the 5mC proficient MEFp (gray) and 5mC deficient MEFpm (yellow). The mean value for the non-barrier (blue) and barrier (magenta) compartments were calculated for each protein in each cell line and statistically compared the barrier one (magenta line) and the overall dips (black lines). n.s.: non significant; ***: p-value < 0.0001. E. Percentage of barrier compartments found in MOCHA-FRAP for the 5mC proficient MEFp and 5mC deficient MEFpm for Mbd1, Mbd2 and MeCP2. Blue box: non-barrier compartments, orange box: barrier compartments. The values on each box represent the mean value of the individual dips for each type of compartment. F. Average MOCHA-FRAP curves from HP1beta on the H3K9me3 proficient MEFw8 (black and gray) and H3K9me3 deficient MEFD5 (dark red). Each compartment was analyzed alone to determine the belonging to barrier or not barrier and then averaged. Lines represent the average of all compartments within the same category and shades the 95% confidence interval. G. Violin plot of the dips calculated for HP1beta on the H3K9me3 proficient MEFw8 (black and gray) and H3K9me3 deficient MEFD5 (dark red). The mean value for the non-barrier (blue) and barrier (magenta) compartments were calculated for each protein in each cell line and statistically compared the barrier one (magenta line) and the overall dips (black lines). n.s.: non significant; *: p-value < 0.05. H. Percentage of barrier compartments found in MOCHA-FRAP for the H3K9me3 proficient MEFw8 and H3K9me3 deficient MEFD5 for HP1beta. Blue box: non-barrier compartments, orange box: barrier compartments. The values on each box represent the mean value of the individual dips for each type of compartment. The number of cells, replicates and the concrete values with error are provided in the Table S8.

As the most predominant barrier compartments were found in Mbds and HP1beta, we tested the effect of the ligand by using mouse embryonic fibroblast cell lines with low to very low levels of the ligands and compare them with their wild type counterparts (Figure 3B): in the case of 5mC, the MEFpm (Lande-Diner *et al*., 2007), knock-out for Dnmt1, the DNA methyltransferase responsible of the maintenance of 5mC upon DNA replication, and for H3K9me3, the MEFD5, knock-out for the two isoforms of the Suv39 methyltransferases (Peters *et al*., 2001), responsible of the methylation of the lysine 9 of histone H3. As Dnmt1 is an essential gene in somatic cells, P53 needs to be also knocked out and therefore is compared to a cell line that also has P53 mutated (MEFp). We screened the heterochromatin compartments with MOCHA-FRAP in these cell lines. The analysis (Table S8) showed that: i) the dip of the barrier compartment generated by Mbds remained unchanged in absence of 5mC in both the average curves (Figure 3C) and the mean of the individual dips (Figure 3D), but MeCP2 showed an overall reduced dip compared to the 5mC proficient cells; ii) HP1beta barrier compartments dip were reduced in absence of H3K9me3, as shown for the average dip (Figure 3F) and the individual dips (Figure 3G); iii) the percentage of compartments showing barriers was slightly reduced for Mbd1, unchanged for Mbd2 and severely reduced for MeCP2 (Figure 3E) in the 5mC deficient cells; iv) the percentage of HP1beta was unchanged in the H3K9me3 deficient cells compared to the proficient (Figure 3H). The results of the 5mC deficient cells is another indication of the differences between *in vitro* LLPS formation and the *in cellulo* behavior of barrier formation, as 5mC *in vitro* promotes Mbd2 LLPS (Zhang *et al*., 2025) but limits LLPS for MeCP2 (Zhang *et al*., 2022; Zhang *et al*., 2023), but in our screening Mbd2 remains mainly unaffected while MeCP2 reduces the ability to form barriers. However, this reduction might be explained by the clustering of barrier compartments, reducing their number (Zhang *et al*., 2022; Brero *et al*., 2005). In contrast, HP1beta is directly regulated by its ligand both *in vitro (Qin, Stengl, et al., 2021)* and in our results.

### 5mC distribution determines the ability of Mbd2 and MeCP2 to form barriers in heterochromatin

We then wondered if we could explain the differences seen between the cell lines as a consequence of a different distribution of the ligands. When analyzing the total levels of the ligands (Table S9), we observed that the levels of 5mC were similar in all three fibroblasts in mouse cells, whereas in C2C12 the levels were lower and in NSC were higher (Figure 4A). The levels of H3K9me3 showed a similar tendency, although there were some differences also between the three fibroblast cell lines (Figure 4C). Analyzing the human cells, hTERT-RPE1 had similar levels to those of NSC for both 5mC (Figure 4B) and H3K9me3 (Figure 4D), while NDF had similar 5mC levels (Figure 4B) and higher H3K9me3 (Figure 4D) than the hTERT-RPE1. We could not find any relation of ligand levels in the different cell lines with barrier formation of heterochromatin of protein binding those ligands.

**Figure 4.**
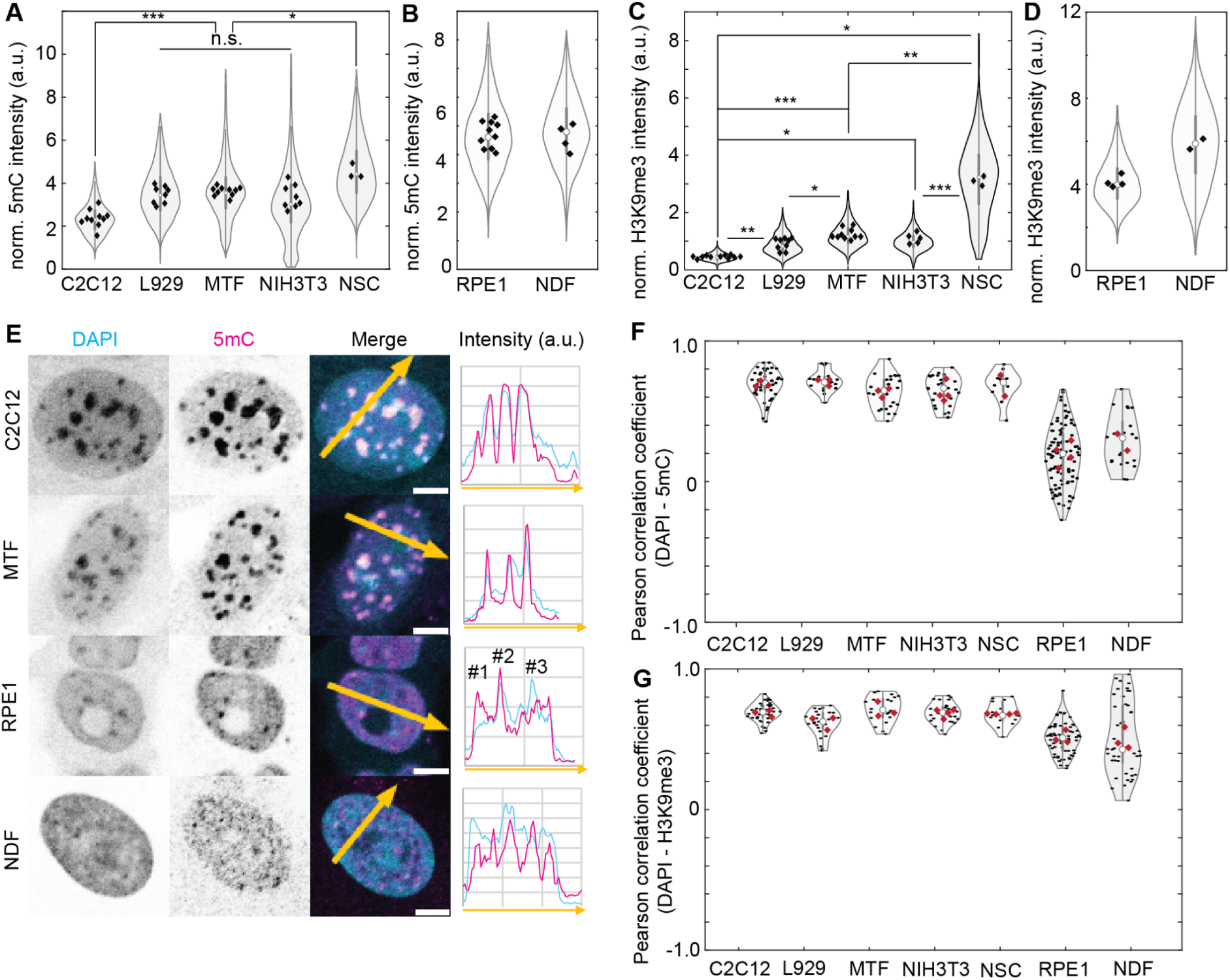
The distribution of the ligands, rather than its levels, explain the differences observed in the barrier formation on the Mbds. A-B. Levels of 5mC normalized to the unit of DNA (DAPI) for mouse (A) and human (B) cells. The violin plots show the distribution for the whole population and the diamonds the mean value for each individual replicates. Cells were fixed with formaldehyde, permeabilized with Triton X-100, treated with RNAseA (to remove the RNA, which can also contain 5mC) and DNaseI (to expose the 5mC) and then incubated with anti-5mC antibody. C-D. Levels of H3K9me3 normalized to the unit of DNA (DAPI) for mouse (C) and human (D) cells. The violin plots show the distribution for the whole population and the diamonds the mean value for each individual replicates. Cells were fixed with formaldehyde, permeabilized with Triton X-100 and incubated with anti-H3K9me3 antibody. The number of cells, mean and 95% confidence interval for each individual replicate of the panels A-D are provided in Table S9. The table also includes the statistics (Hedges G and Welch p-value) to compare the different conditions. E. Examples of cells stained with 5mC. The graph is the intensity over the arrow that is shown in the merge (average of the 4 pixel in each point of the line). In hTERT-RPE1 cells, the peaks correspond to the three possibilities observed in the cells: #1 is a 5mC focus not related with heterochromatin, #2 is a 5mC focus within heterochromatin and #3 is a cluster of 5mC focus near the heterochromatin. F-G. Pearson correlation coefficient calculated for different replicates for 5mC versus DAPI (F) or H3K9me3 versus DAPI (G) for each cell line used in the study. The values of each replicate, as well as the number of cells, are provided in Table S10.

However, we found a striking difference between the mouse and human cell lines regarding 5mC distribution (Figure 4E). Whereas in mouse 5mC perfectly aligned with the DAPI intensity in all cell lines, this was not the case for the human cells. When we analyzed the position of the foci related to the perinucleolar heterochromatin compartments seen in DAPI, we found three different kind of foci as shown in the hTERT-RPE1 cell example (Figure 4E, middle down panel): i) foci located out of the perinucleolar compartments (marked as #1); ii) foci located within the heterochromatin (marked as #2); and iii) foci located in the proximity of the perinucleolar compartments (marked as #3). The rarest ones were the ones on heterochromatin. This was not the case for the NDF cells, where foci were located within the heterochromatin as well as in close proximity, even if it was not covering the whole compartment as it happened in the mouse cells. In contrast, H3K9me3 foci covered the heterochromatin compartments in all cell lines. We quantified this phenomenon using Pearson correlation coefficient (PCC) on the middle plane of cells immunostained with anti-H3K9me3 or anti-5mC antibodies and comparing the staining with the DAPI signal (Figure 4F-G). In the case of 5mC, all mouse cells showed high and comparable values of correlation, between 0.632 and 0.708 (Table S10), while hTERT-RPE1 cells lost it completely with an average of 0.198 (Figure 4F). NDF had an intermediate correlation but, more importantly, the variance between cells was less, therefore having higher probability of having at least some correlation compared to the hTERT-RPE1 cells. This was not the case for H3K9me3 (Figure 4G), were the values in mouse were in the same range as for 5mC (0.609 to 0.708), but the human cells were having only slightly reduced (0.515) (Table S10), likely due to the intensity differences in DAPI as a result of the increased AT-rich regions in mouse heterochromatin. Indeed, only in hTERT-RPE1 cells there are significant differences between the PCC of 5mC against DAPI and H3K9me3 against DAPI, indicating the loss of 5mC in the heterochromatin compartments (Table S10). Therefore, we concluded that it was the distribution of 5mC, and not its total levels, the decisive parameter that explained the changes observed in the barrier formation on the Mbd proteins.

The fact that the 5mC distribution is more relevant in humans than in mouse cells implies the presence of other factors that bring the Mbd proteins to the heterochromatin. Mbd1 and MeCP2 have known interactions that help them to be in the mouse heterochromatin, as the CxxC3 domain of Mbd1 binds to unmethylated CpG (Jørgensen *et al*., 2004) and the three AT-hooks, especially AT-hook3 of MeCP2 makes MeCP2 bind preferentially to AT-rich regions (Baker *et al*., 2013; Lyst *et al*., 2016). Both CpG and AT-rich regions are enriched in the major satellite repeats that form the majority of the pericentric heterochromatin in mouse cells. While CpG could be still enriched in the perinucleolar heterochromatin in hTERT-RPE1 and NDF cells, there is no evidence of AT-rich as shown by the lower accumulation of DAPI in these compartments. In addition to the AT-hook, MeCP2 recruitment to heterochromatin in mouse is also dependent on major satellite RNA (Fioriniello *et al*., 2020), and this could also be the mechanism Mbd2 utilizes due to the presence of RG domains that have been described to interact with RNA in other proteins (Doron-Mandel *et al*., 2021; Ozdilek *et al*., 2017). The α-satellite RNA in human cells has been shown to interact and stabilize Suv39H1, therefore contributing to heterochromatin formation by H3K9me3 (Johnson *et al*., 2017) but there is no evidence of recruitment of Mbds. The combination of unmethylated CpGs, AT-rich regions and RNA produce the different binding from the Mbds to the heterochromatin and therefore could affect their ability of barrier formation.

### Heterochromatin compartments compete for the free pool of nucleoplasmic protein to form barrier compartments

Based on the data we collected, we defined some assumptions to model the behavior of the proteins of interest (POIs) that are susceptible to barrier formation of heterochromatin (Figure 5A). These assumptions were the following: i) chromatin is represented as a polymer containing a heterochromatic region; ii) heterochromatic regions contain anchors that can self-interact and form clustered structures; iii) POIs can establish multivalent interactions with the heterochromatic binding sites and with themselves with similar affinity. Note that anchor means in this model any sequence or ligand that would help the POI to enrich, including H3K9me3/5mC, RNA, AT-regions, CpG sequences or other protein components that could potentially recruit specifically the POI. We settled on three initial conditions, with the only difference between them being the levels of the POI, as low, medium or high, and studied how heterochromatin organization and barrier formation changed among these conditions (Figure 5B). At low concentrations (Figure 5B, upper panels), the POI occupied the binding sites in heterochromatin without recruiting additional protein layers, indicating the absence of a multivalent network and any barrier compartment. At medium concentrations (Figure 5B, middle panels), saturation of binding sites by POI binding is followed by the nucleation of a multivalent network of POIs. Since heterochromatic regions are tethered to the simulation box, the cluster can only grow through diffusion of molecules from smaller to larger structures. This competition between heterochromatic foci leads to co-existence of phase-separated compartments that display a barrier with compartments where the POIs are bound without forming a multivalent network. Finally, when high concentrations of the POI were used (Figure 5B, lower panels), the competition was unnecessary as there was enough free pool of POI to allow all compartments to become barrier compartments.

**Figure 5.**
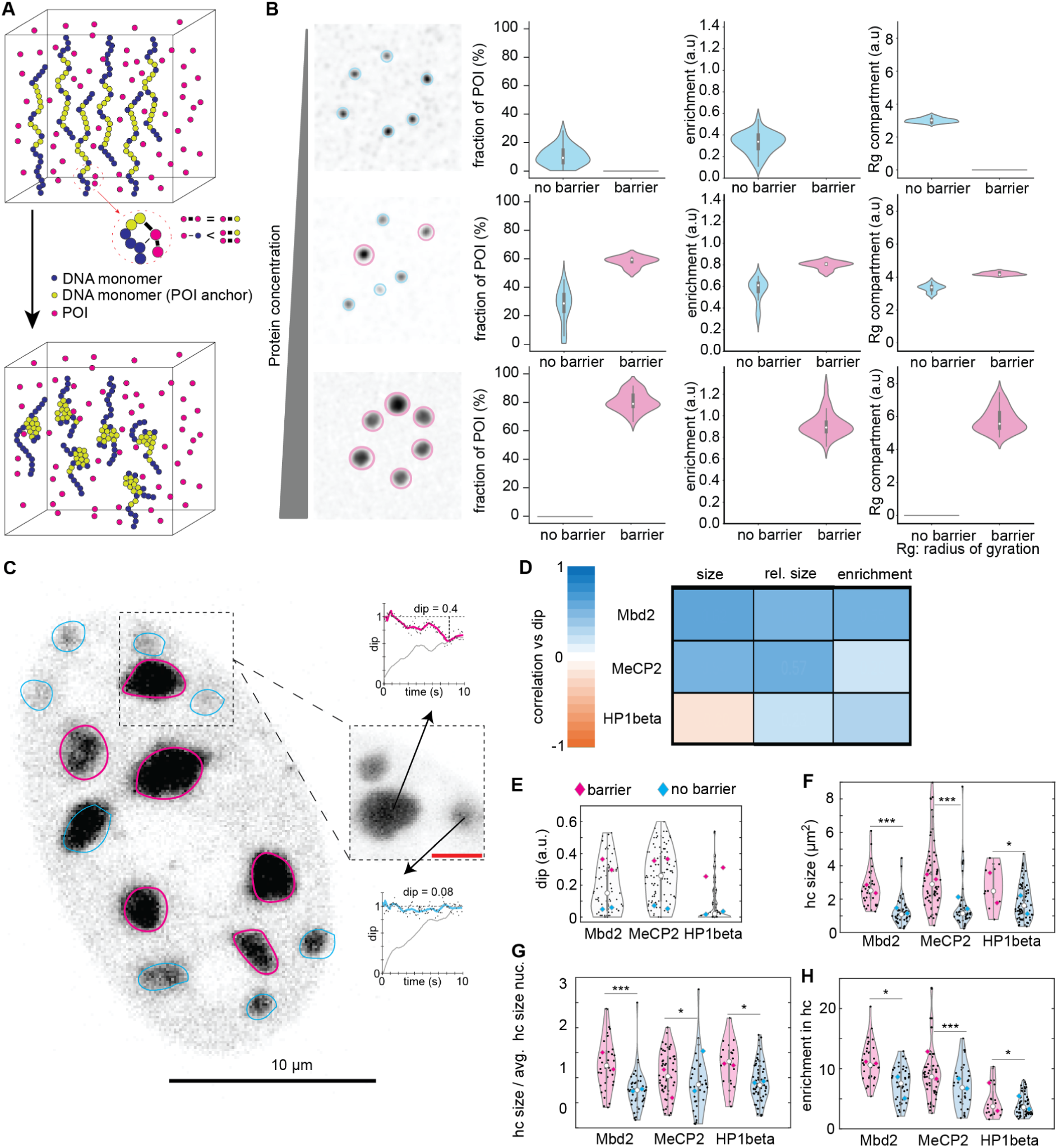
A compartment competition model to generate barrier compartments. A. Initial conditions for the model. The DNA is defined as a polymer, with the chromatin monomers representing a unit of DNA that can be enriched (yellow) or not (blue) in anchors for the protein of interest (POI). These anchors can be the ligand for the POI (5mC or H3K9me3) but also any other factor on the DNA that could bring the POI to the heterochromatin (i.e. unmethylated CpG for Mbd1). The POI (magenta) can be given at different concentrations. The self-interaction of the POI that allows the LLPS and the barrier formation has the same strength as the interaction of the POI with the chromatin monomer harboring the binding sites. Clustering of the heterochromatin regions is performed before establishing chromatin-POI interactions. B. Results of running the model with low (upper panel), medium (middle panel) or high (lower panel) POI concentration as starting conditions. Left column: visualization of the POI accumulating on the heterochromatin compartments as result of multivalent POI interactions; middle column: enrichment of the POI in the barrier or non barrier compartments; right column: size of the non barrier or barrier compartments. C. Representative image of the MOCHA-FRAP experiment of all compartments in one cell, using C2C12 and L929. Every compartment was analyzed with MOCHA-FRAP in its middle point (as shown in the zoomed box, where two of the three compartments are out of focus in the overview but in the middle plane in the zoom). The compartments were assigned the category barrier or non barrier based on the dip and used for the further calculations. D. Correlation of dip with size, relative size (size of the compartment divided by the average size compartment in the cell) or enrichment for the three proteins studied. The data from both C2C12 and L929 cell lines were combined for this plot. E-H. Distribution of the values of dip (E), heterochromatin compartment size (F), relative heterochromatin compartment size (G) and enrichment (H) for barrier and non barrier compartments for each protein. The average values for C2C12 and L929 are separated in each case (diamonds located in the left, C2C12; in the right, L929). A two sided t-test was used to compare the barrier versus the non barrier compartments. *: p-value < 0.05; ***: p-value < 0.0001.

Interestingly, we observed that, in the model, the barrier compartments grow larger, due to the fact that they attracted more neutral polymer in, and were more enriched in the POI than the non-barrier compartments as a consequence of the barrier formation (Figure 5B). We tested this observation on the screening results in mouse and human cells, correlating the dip in the compartment against the size of the compartment (Figure S6A) or versus the enrichment of the protein in the compartment (Figure S6B). While the size correlation worked for most of the proteins studied, the enrichment correlation was more difficult to interpret. We thought that the optimization done in light intensity for each individual compartment might affect the enrichment by changing the nucleoplasm intensity, making these results non-comparable. Therefore, we performed additional MOCHA-FRAP experiments in which (Figure 5C): i) we studied all of the heterochromatin compartments within a cell, each of them in the middle point; ii) we preserved the laser intensities within a cell to have a consistent enrichment data: iii) we analyzed the compartments individually and classified them in barrier or not barrier based on the MOCHA-FRAP dips. We did this experiment with three different proteins, Mbd2, MeCP2 and HP1beta, and two different cell lines, C2C12 and L929. We selected cells ranging from low-middle to high protein levels for each protein of interest. When we analyzed the size (Figure 5D), Mbd2 and MeCP2, as in the screen (Figure S6A), showed a strong correlation with the dip, and the barrier compartments were larger than the non-barrier ones (Figure 5F). In HP1beta, the correlation was even slightly negative (Figure 5D) as it was in the screen but there was a tendency for the barrier compartments to be bigger than the non-barrier ones (Figure 5F). However, when we analyzed the size of the compartments normalized to the average on the cell, the differences were more clear in both the correlation with the dip (Figure 5D) and the differences in size between the barrier and non-barrier compartments (Figure 5G). Finally, the enrichment of the proteins was directly correlating with the dip for all proteins (Figure 5E) and was significantly higher in barrier compartments (Figure 5H).

Based on the model, we could identify the enrichment of the proteins as the most relevant factor in the formation of barrier compartments. In our screen, the loss of barrier compartments in human cell lines came together with a reduction in the overall enrichment of the protein of interest compared to the mouse cell lines. In Mbd2, mouse cells have an average enrichment of 9.2 in heterochromatin and 37.8% of barrier compartments, and this enrichment drops to 3.7 in hTERT-RPE1 cells with a total loss of the barrier compartments and to 5.9 in NDF where there is still 11.1% of barrier compartments. In MeCP2, the reduction of the enrichment is similar between the two cell lines, from an average enrichment of 8.4 in mouse cells to 2.0 and 2.6 in hTERT-RPE1 and NDF cells, respectively, and the percentages of barrier compartments also drop similarly from 50.2% average in mouse to 13.5% and 18.6% in hTERT-RPE1 and NDF, respectively. In the case of HP1beta, the reduction of the enrichment is not as drastic, from 5.8 average in mouse to 5.4 and 5.0, but in this case the increase of HP1alpha in heterochromatin might be limiting the access to H3K9me3, which is necessary for HP1beta barrier formation (Qin, Stengl, *et al*., 2021) as shown in the H3K9me3 deficient mutants (Figure 3G).

### Cells have multiple ways to regulate barrier formation on heterochromatin

Based on our results, we defined a working model for the formation of barrier compartments of heterochromatin for a specific POI that is regulated at four different levels: i) the heterochromatic environment; ii) the concentration of the POI; iii) the competition among compartments for the nucleoplasm POI; and iv) the multivalence of the interactions of the POI.

The heterochromatic environment defines how well the POI can be retained in heterochromatin. It is defined by the high affinity ligands of the POI (5mC for MBDs and H3K9me3 for HP1s), but also for other anchor points, defined as other proteins, DNA or RNA that interact with the POI and allow a transient retention of it in the heterochromatin, or by other proteins, DNA or RNA that compete with the POI for those anchor points.

The POI starts to accumulate in heterochromatin thanks to these high affinity or transient interactions, which allows the self interaction of the POI in the heterochromatin. However, the formation of the barrier compartments requires additional POI recruitment/retention in heterochromatin through self-interactions. Therefore, the concentration of the protein in the nucleus should be sufficient to fill the anchor points in heterochromatin and leave a nucleoplasm pool that can potentially be recruited to the heterochromatin until a threshold is reached where the POI self-interaction allows the formation of a barrier.

Once one compartment forms a barrier, there is a competition between compartments for the free POI in the nucleoplasm. The concentration of the protein in the nucleus will determine how strong this competition is, as increasing concentration will assure that more compartments can achieve the required threshold to form the barrier compartment.

Finally, the multivalence of the POI will determine the ability of compartments to grow, as the same number of POI within a barrier compartment will be able to take more POI from the nucleoplasm pool the more multivalency it has. This is potentially the reason why barrier and non-barrier compartments differ in size and enrichment from MeCP2/Mbd2 compared to HP1beta. In this case, the differences in the intrinsically disorder regions between proteins is also relevant for that purpose.

These regulatory steps explain the disparate results obtained in different studies regarding the same protein. If we take the example of HP1alpha, we observed that it was not able to form a barrier in adult stem cells (C2C12 and NSC) as well as non-stem cells such as fibroblasts (L929, MTF and NIH-3T3) (Figure 1). We could reproduce the results of Erdel et al. (Erdel *et al*., 2020) in the same system (NIH-3T3). However, another study that was done in embryonic stem cells (Gaurav *et al*., 2025) showed that HP1alpha was capable of form barriers in this system in the presence of KAP1, which enhanced the LLPS abilities of HP1alpha *in vitro*. In this case, KAP1 would act as an anchor point. On the other hand, the presence of other HP1s reduces HP1alpha ability to undergo LLPS (Bensaha *et al*., 2025) by competing for the anchor points and make the transient accumulation more unstable. Additional studies showed that blocking weak interactions with 1,6-hexanediol (Strom *et al*., 2017) or the dimerization ability (Larson *et al*., 2017) of HP1alpha reduced the enrichment and the presence of larger heterochromatin compartments, respectively. Lastly, orthologues of HP1alpha have been shown to be more prone to LLPS than the mouse HP1alpha, probably due to changes in its intrinsically disorder domains (Bensaha *et al*., 2025), therefore suggesting that the inherent abilities of the protein also influence the formation of barrier compartments.

Previous work averaged all compartments together and, thus, could not identify barrier characteristics provided by only a subset of those compartments. Our present analysis demonstrated that the latter can range from 10 to 70% of all compartments. Future studies should take in consideration the individuality of the compartments and categorize them accordingly.

In summary, our findings and heterochromatin compartment competition model resolve the existing controversy and rationalize how in cells heterochromatin compartments form and compete to establish dynamic barriers to the entry and exit of its components.

## Author contributions

Experimental design: H.R. and M.C.C.; MOCHA-FRAP acquisition: M.A., H.R., H.Z. and M.M.; MOCHA-FRAP analysis: H.R.; immunofluorescence staining: M.A. and A.Z.; high-throughput acquisition and analysis: A.Z.; Interactome profiling: W.Q. and H.L.; Modelling: F.M. and F.E.; Confocal microscopy acquisition and analysis: H.R.; Funding acquisitions: M.C.C. and H.L.; Visualization: H.R.; Writing - original draft: H.R.; Writing - review and editing: H.R. and M.C.C.

## Acknowledgements

We thank Anne Lehmkuhl for technical support in the maintenance of the different cell lines, and Katerina Gurova for providing the human NDF cell line.

## Funding

This work was funded by the Deutsche Forschungsgemeinschaft (DFG, German Research Foundation) grants CA 198/19-1 project number 522122731 to M.C.C. and LE 721/18-1 project number 425470807 to H.L..

## Conflict of interest

The authors declare no conflict of interests.

## Renewal materials, data and code availability

The raw data and analyses, including the self-written code that support the findings of the study are available in the TUdatalib repository with the identifier https://doi.org/10.48328/tudatalib-2252. Renewal materials are available through the corresponding author upon reasonable request.

## Supplementary figures

**Figure S1.**
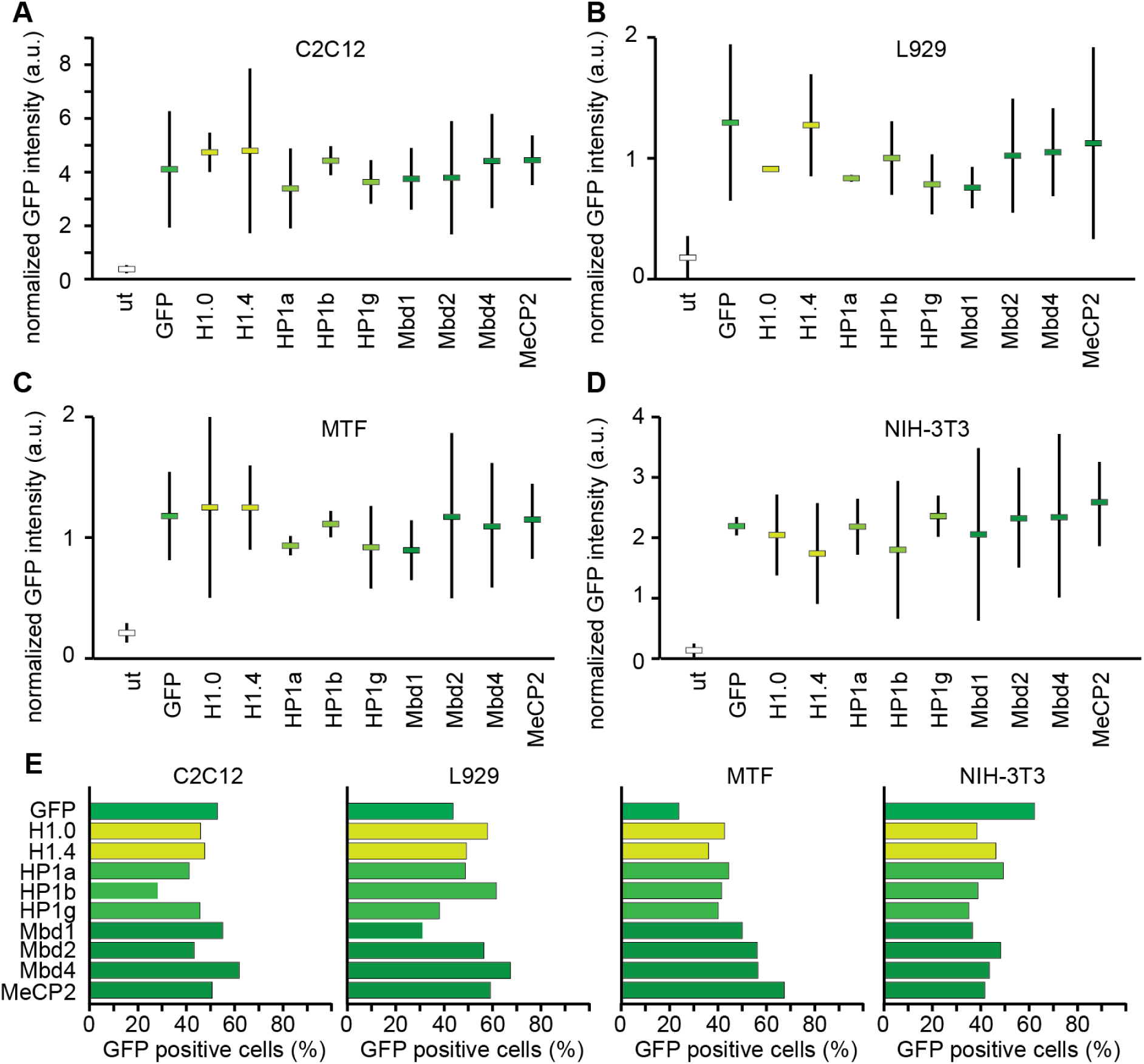
Protein level in transfected cells is similar among the plasmids used. A-D. Average expression levels for positive cells for the proteins used in the study in C2C12 (A), L929 (B), MTF (C) and NIH-3T3 (D). Cells were transfected, fixed, counterstained with DAPI and the levels of GFP in the nucleus was quantified using high throughput microscopy. The analysis pipeline segmented the cell based on the nucleus properties (size and roundness) using the DAPI (DNA) signal and then calculated the total intensity in the GFP channel and normalized it to the total intensity in DAPI to avoid cell cycle variations. Positive cells were defined as normalized GFP intensity higher than the 95 percentile of the untransfected sample. The graph represents the mean and 95% confidence interval of at least two independent replicates. E. Percentage of positive cells obtained for each protein of interest in each cell line.

**Figure S2.**
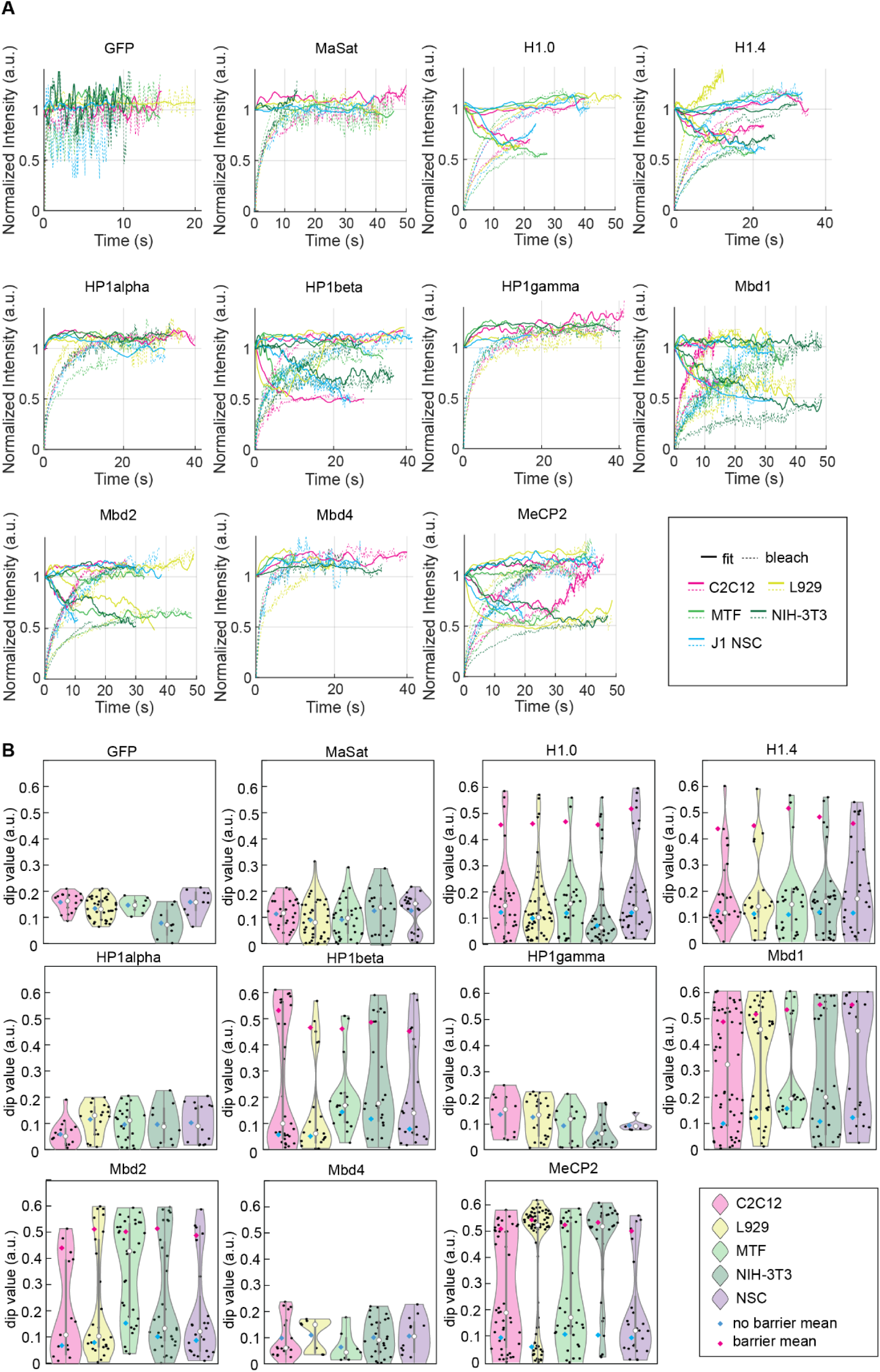
MOCHA-FRAP screening in mouse cell lines. A. Average graphs for the barrier and non-barrier compartments in each cell line. For better visualization, only the Savitzky-Golay fit from the non-bleached half (that is used to calculate the dip) and the average bleach curve are shown for each cell line. The complete graphs with error and the average non-bleach curve can be found in the raw data 1 in TUdatalib. B. Violin plot with the dip of the individual compartments used for calculating the average graphs. Black dots represent the individual dips, blue diamonds the mean of the non-barrier compartments, magenta diamonds the mean of the barrier compartments, white dot is the median of all data, gray box represent the 25 to 75 percentile and the gray line the standard deviation of the data. The number of cells and biological replicates, as well as the concrete values shown in the graphs are provided in the Table S5.

**Figure S3.**
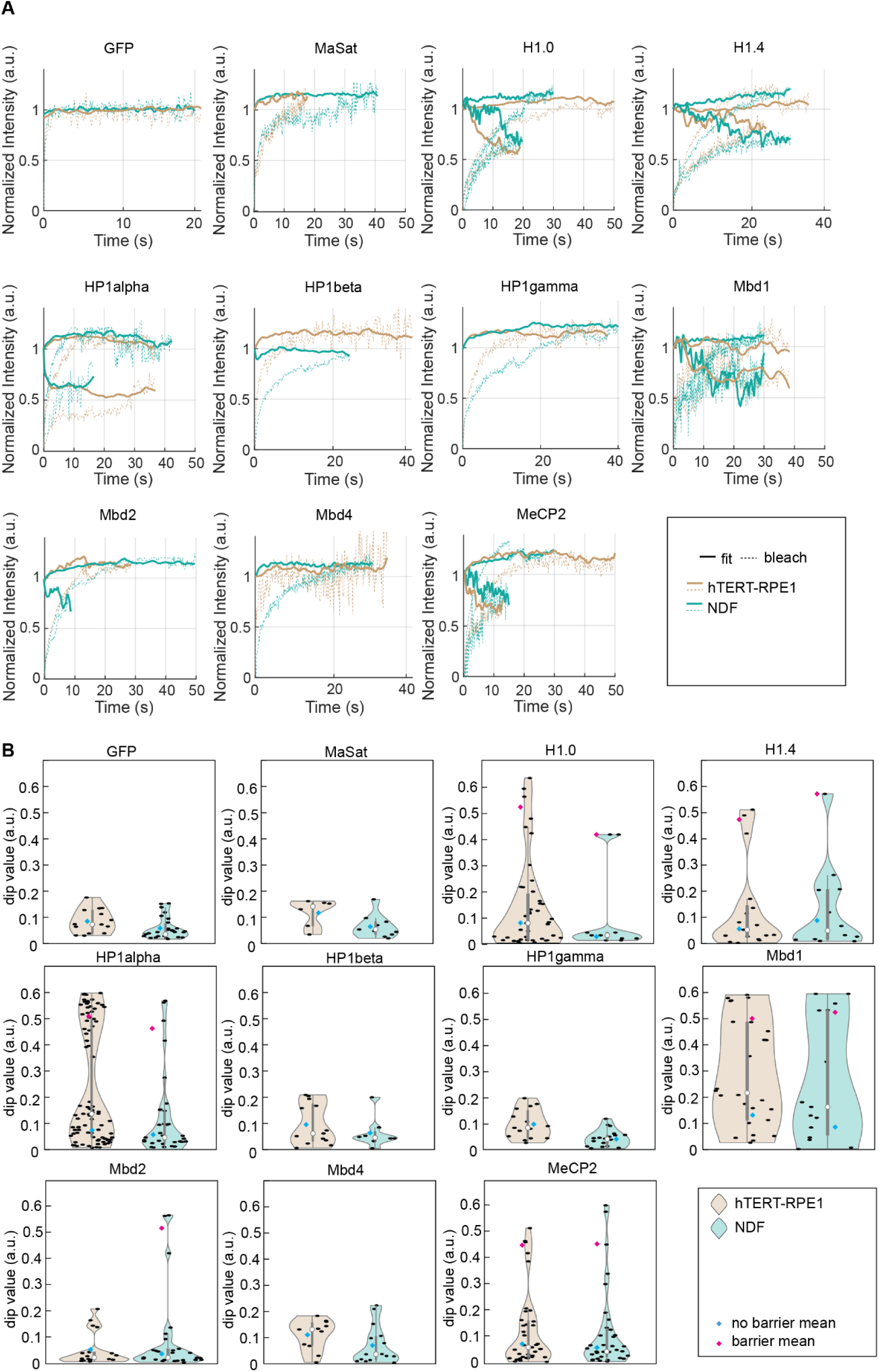
MOCHA-FRAP screening in human cell lines. A. Average graphs for the barrier and non-barrier compartments in each cell line. For better visualization, only the Savitzky-Golay fit from the non-bleached half (that is used to calculate the dip) and the average bleach curve are shown for each cell line. The complete graphs with error and the average non-bleach curve can be found in the raw data 1 in TUdatalib. B. Violin plot with the dip of the individual compartments used for calculating the average graphs. Black dots represent the individual dips, blue diamonds the mean of the non-barrier compartments, magenta diamonds the mean of the barrier compartments, white dot is the median of all data, gray box represent the interquartile distance (25 to 75 percentile) and the gray line the standard deviation of the data. The number of cells and biological replicates, as well as the concrete values shown in the graphs are provided in Table S6.

**Figure S4.**
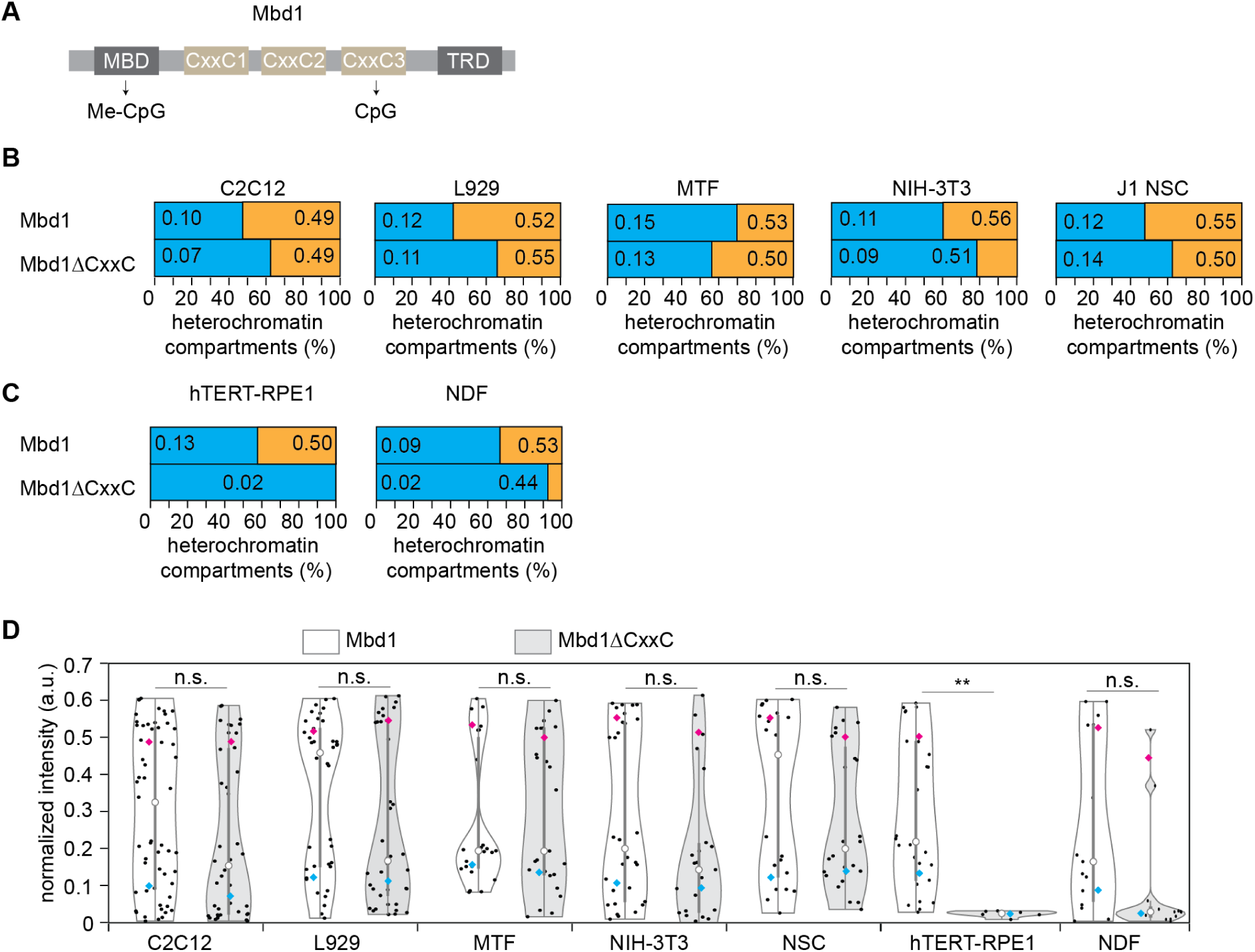
Mbd1ΔCxxC ability to form barrier compartments is decreased in human cells. A. Overview of the main domains of the Mbd1 protein. The MBD domain is responsible for the binding to 5mC. CxxC1 and CxxC2 are not fully functional, so only CxxC3 is responsible for binding to unmethylated CpG. CxxC3 is not present in the short isoform Mbd1ΔCxxC. B-C. Barrier compartment formation analyzed by MOCHA-FRAP of Mbd1ΔCxxC in the mouse (B) and human (C) cell lines used for the screening. The Mbd1 results from figures 1K and 2I were added for comparison. Blue represents non-barrier compartments and orange barrier compartments. When both are present, left hand side numbers represent the average dip from the individual non-barrier compartments and right hand side numbers the average dip from the individual barrier compartments. D. Violin plot representing the individual points used to calculate the percentage and the mean values shown in B and C. A double sided t-test compared the Mbd1 versus Mbd1ΔCxxC dip. n.s.: non significant (p-value > 0.05); **: p-value < 0.001

**Figure S5.**
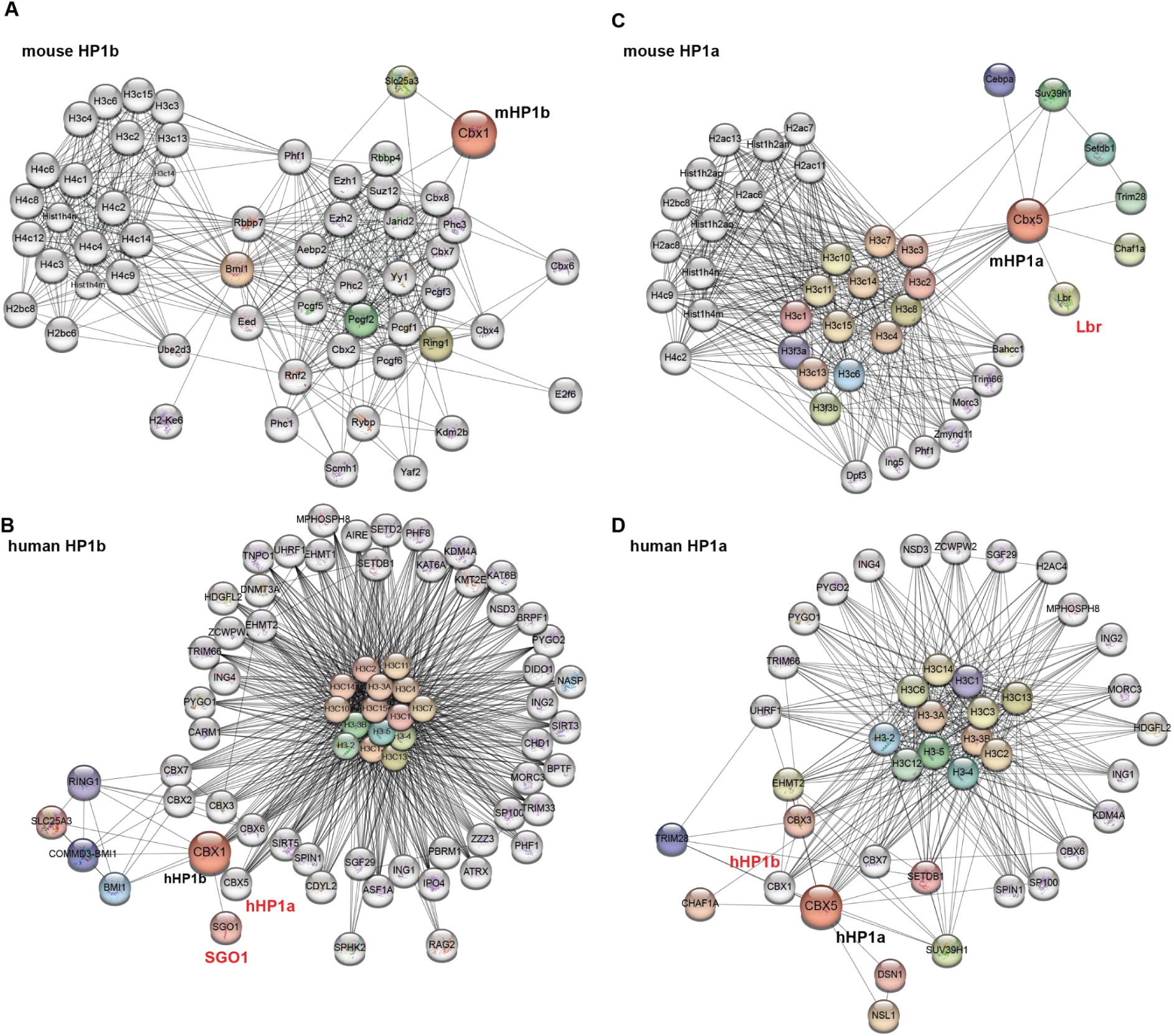
HP1beta and HP1alpha interactome in mouse and human cells based on the STRING database. Relevant changes have been marked in red. Colored nodes represent the query proteins and proteins closely associated with them, whereas white nodes indicate secondary interactors or proteins automatically added by STRING. A-B. Interactome of HP1beta in mouse (A) and human (B) cells. C-D. Interactome of HP1alpha in mouse (C) and human (D) cells.

**Figure S6.**
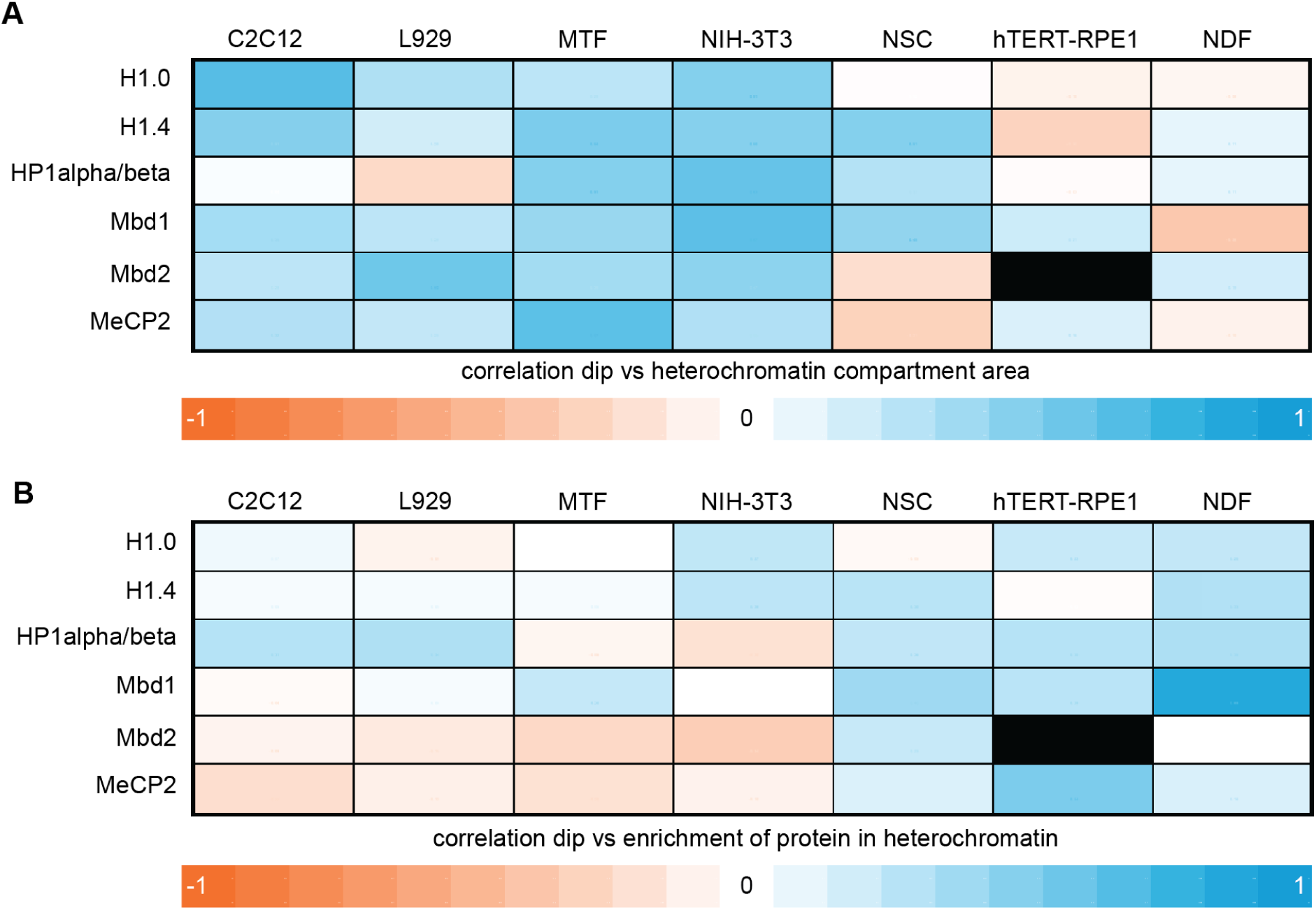
Correlation of the screenings in mouse and human cells to the characteristics of heterochromatin. A. Heat map correlation of the dip for the protein of interest in heterochromatin compartments with the area of the compartments. B. Heat map correlation of the dip for the protein of interest in heterochromatin compartments with the enrichment of the protein in the compartments. The values of the mean area and enrichment, as well as the values of the correlation of these parameters with the dip, are provided in Table S7.

## Supplementary tables

**Supplementary Table S1:** Cell line characteristics and transfection.

| Name | Species | Type | Genotype | AMAXA program | Reference |
| --- | --- | --- | --- | --- | --- |
| C2C12 | *Mus musculus* | myoblast | wild type | B-032 | (Yaffe and Saxel, 1977) |
| J1 NSC | *Mus musculus* | neural stem cell | wild type | A-033 | (Romero *et al.*, 2025) |
| L929 | *Mus musculus* | connective tissue fibroblast | wild type | T-020 | (Sanford *et al.*, 1948) |
| MEF D5 | *Mus musculus* | embryonic fibroblast | *Suv39h1<sup>-/-</sup>*<br>*Suv39h2<sup>-/-</sup>* | A-023 | (Peters *et al.*, 2001) |
| MEF-P | *Mus musculus* | embryonic fibroblast | *P53<sup>-/-</sup>* | A-023 | (Lande-Diner *et al.*, 2007) |
| MEF-PM | *Mus musculus* | embryonic fibroblast | *P53<sup>-/-</sup> Dnmt1<sup>-/-</sup>* | A-023 | (Lande-Diner *et al.*, 2007) |
| MEF W8 | *Mus musculus* | embryonic fibroblast | wild type | A-023 | (Peters *et al.*, 2001) |
| MTF | *Mus musculus* | tail fibroblast | wild type | A-023 | (Guy *et al.*, 2001) |
| NIH-3T3 | *Mus musculus* | embryonic fibroblast | wild type | A-023 | (Jainchill *et al.*, 1969) |
| hTERT RPE-1 | *Homo sapiens* | hTERT immortalized retinal pigment epithelial cell | wild type | X-001 | (Bodnar *et al.*, 1998) |
| hNDF | *Homo sapiens* | Neonatal dermal fibroblasts | wild type | U-023 | (Leonova *et al.*, 2018) |

**Supplementary Table S2:** Plasmid characteristics.

| Name | pc number* | Fluorophore | Gene species | Promoter | Addgene | Reference |
| --- | --- | --- | --- | --- | --- | --- |
| pEGFP-N1 | 0713 | GFP | *Aequorea victoria* | CMV |  | Clontech |
| pMaSat-GFP | 1803 | GFP | Synthetic | CMV |  | (Lindhout *et al.*, 2007) |
| pMaSat-mRFP | 2063 | mRFP | Synthetic | CMV | 201695 | (Herce *et al.*, 2017) |
| pGFP-H1.0 | 5157 | GFP | *Homo sapiens* | CMV |  | (Th'ng *et al.*, 2005) |
| pGFP-H1.4 | 2378 | GFP | *Homo sapiens* | CMV |  | (Th'ng *et al.*, 2005) |
| pGFP-HP1alpha | 1174 | GFP | *Homo sapiens* | CMV | 17652 | (Cheutin *et al.*, 2003) |
| pdsRed-HP1alpha | 1224 | dsRed | *Homo sapiens* | CMV |  | (Hayakawa *et al.*, 2003) |
| pGFP-HP1beta | 1175 | GFP | *Homo sapiens* | CMV | 17651 | (Cheutin *et al.*, 2003) |
| pGFP-HP1gamma | 1176 | GFP | *Homo sapiens* | CMV | 17650 | (Cheutin *et al.*, 2003) |
| pMbd1-GFP | 1191 | GFP | *Mus musculus* | CMV |  | (Hendrich and Bird, 1998) |
| pGmMbd1.3 | 2899 | GFP | *Mus musculus* | CMV | 246850 | (Zhang *et al.*, 2017) |
| peMBD2G | 2399 | GFP | *Mus musculus* | CMV | 211572 | (Becker *et al.*, 2013) |
| pMbd4-GFP | 1194 | GFP | *Mus musculus* | CMV |  | (Hendrich and Bird, 1998) |
| pEG-MeCP2 | 1208 | GFP | *Homo sapiens* | CMV |  | (Kudo *et al.*, 2003) |
| pHTHMeCP2 | 3972 | Halotag | *Homo sapiens* | CMV | 227639 | (Schmidt *et al.*, 2024) |

\* pc: plasmid collection

**Supplementary Table S3:**
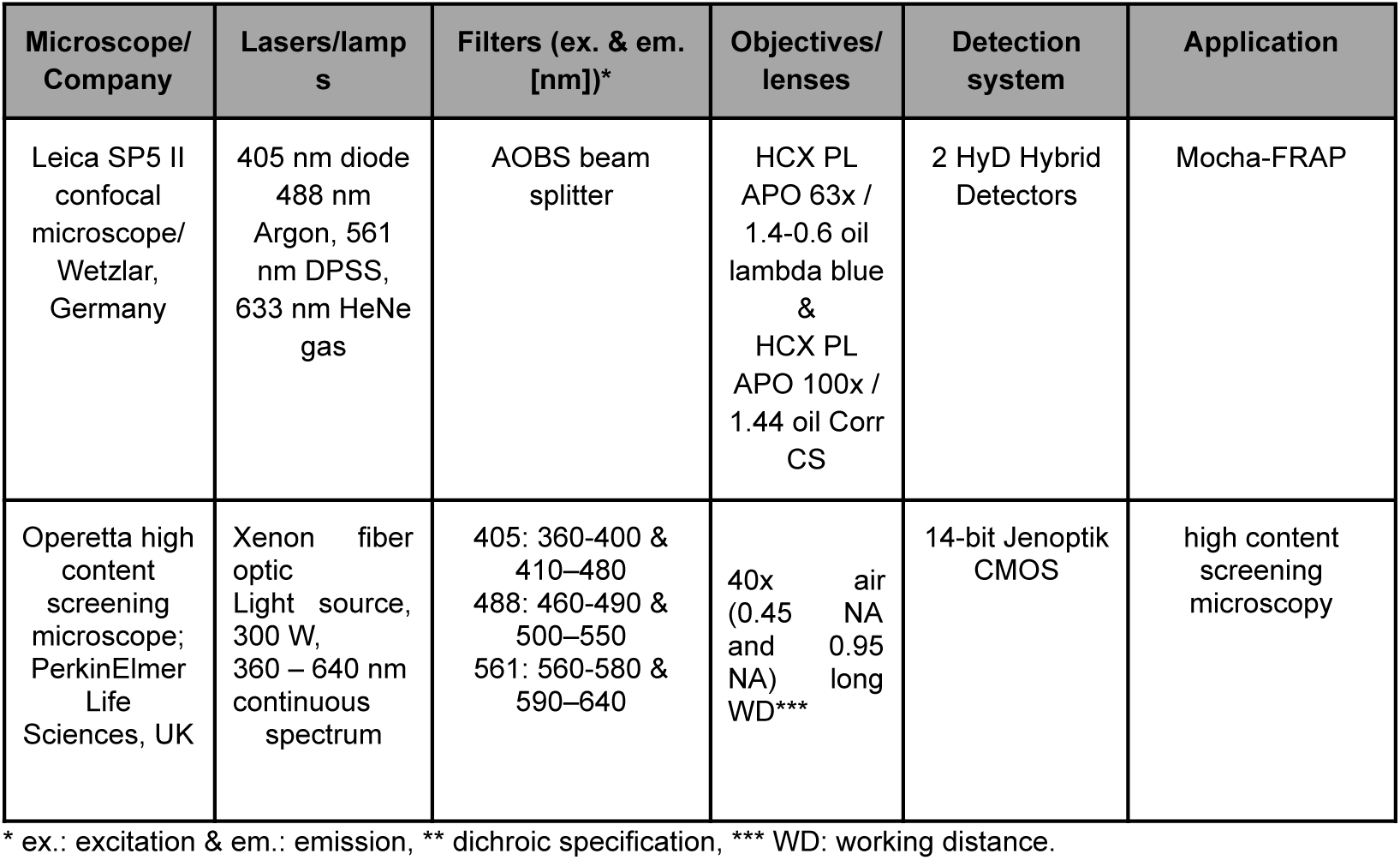
Imaging systems characteristics.

**Supplementary Table S4:** Primary and secondary antibody characteristics.

| Reactivity | Host | Dilution | Cat # | Company/Reference |
| --- | --- | --- | --- | --- |
| α-histone H3 K9 trimethylation | rabbit | 1:100 | 39161 | Active Motif, La Hulpe, Belgium |
| α-5-methylcytosine | mouse | 1:250 | Clone 32E2 | (Weichmann *et al.*, 2020) |
| α-mouse IgG Cy3-conjugated | donkey | 1:400 | JIM-711-16<br>5-152 | The Jackson Laboratory, Bar<br>Harbor, ME, USA |
| anti rabbit IgG Cy3-conjugated | goat | 1:250 | R37117 | Thermo Fisher Scientific, Waltham,<br>MA, USA |

**Supplementary Table S5.** Dip values (in arbitrary units) from Figure 1 and S2, provided as mean ± 95% confidence interval.

| POI | cell line | replicates | condition | dip (individual) | dip (average) | n (%) |
| --- | --- | --- | --- | --- | --- | --- |
| GFP | C2C12 | 3 | no barrier | 0.16 ± 0.02 | 0.11 ± 0.07 | 19 (100) |
| GFP | L929 | 4 | no barrier | 0.14 ± 0.01 | 0.06 ± 0.06 | 36 (100) |
| GFP | MTF | 2 | no barrier | 0.15 ± 0.02 | 0.10 ± 0.11 | 9 (100) |
| GFP | NIH3T3 | 1 | no barrier | 0.09 ± 0.05 | 0.22 ± 0.44 | 8 (100) |
| GFP | J1 NSC | 1 | no barrier | 0.17 ± 0.04 | 0.01 ± 0.13 | 9 (100) |
| MaSat | C2C12 | 5 | no barrier | 0.12 ± 0.02 | 0.00 ± 0.06 | 27 (100) |
| MaSat | L929 | 4 | no barrier | 0.10 ± 0.02 | 0.05 ± 0.14 | 38 (100) |
| MaSat | MTF | 4 | no barrier | 0.10 ± 0.03 | 0.06 ± 0.12 | 32 (100) |
| MaSat | NIH3T3 | 2 | no barrier | 0.13 ± 0.04 | 0.00 ± 0.10 | 18 (100) |
| MaSat | J1 NSC | 3 | no barrier | 0.13 ± 0.03 | 0.01 ± 0.17 | 21 (100) |
| H1.0 | C2C12 | 3 | barrier | 0.45 ± 0.20 | 0.34 ± 0.17 | 5 (13.9) |
| H1.0 | C2C12 | | no barrier | 0.12 ± 0.03 | 0.06 ± 0.03 | 31 (86.1) |
| H1.0 | L929 | 3 | barrier | 0.46 ± 0.12 | 0.39 ± 0.17 | 6 (11.8) |
| H1.0 | L929 | | no barrier | 0.09 ± 0.02 | 0.01 ± 0.03 | 45 (88.2) |
| H1.0 | MTF | 3 | barrier | 0.46 ± 0.32 | 0.47 ± 0.22 | 3 (10.7) |
| H1.0 | MTF | | no barrier | 0.11 ± 0.03 | 0.05 ± 0.03 | 25 (89.3) |
| H1.0 | NIH3T3 | 3 | barrier | 0.46 ± 0.08 | 0.41 ± 0.03 | 6 (17.1) |
| H1.0 | NIH3T3 | | no barrier | 0.07 ± 0.02 | 0.01 ± 0.04 | 29 (82.9) |
| H1.0 | J1 NSC | 3 | barrier | 0.51 ± 0.06 | 0.39 ± 0.09 | 7 (20.6) |
| H1.0 | J1 NSC | | no barrier | 0.11 ± 0.03 | 0.04 ± 0.08 | 27 (79.4) |
| H1.4 | C2C12 | 3 | barrier | 0.43 ± 0.12 | 0.31 ± 0.08 | 5 (16.7) |
| H1.4 | C2C12 | | no barrier | 0.12 ± 0.03 | 0.05 ± 0.04 | 25 (83.3) |
| H1.4 | L929 | 3 | barrier | 0.45 ± 0.15 | 0.28 ± 0.20 | 4 (21.1) |
| H1.4 | L929 | | no barrier | 0.11 ± 0.03 | 0.03 ± 0.07 | 15 (78.9) |
| H1.4 | MTF | 3 | barrier | 0.51 ± 0.16 | 0.55 ± 0.32 | 3 (10.7) |
| H1.4 | MTF | | no barrier | 0.11 ± 0.03 | 0.01 ± 0.03 | 25 (89.3) |
| H1.4 | NIH3T3 | 3 | barrier | 0.48 ± 0.08 | 0.33 ± 0.16 | 5 (15.6) |
| H1.4 | NIH3T3 | | no barrier | 0.11 ± 0.03 | 0.01 ± 0.06 | 27 (84.4) |
| H1.4 | J1 NSC | 3 | barrier | 0.45 ± 0.06 | 0.40 ± 0.12 | 8 (27.6) |
| H1.4 | J1 NSC | | no barrier | 0.11 ± 0.04 | 0.01 ± 0.04 | 21 (72.4) |
| HP1 alpha | C2C12 | 2 | no barrier | 0.06 ± 0.03 | 0.05 ± 0.07 | 16 (100) |
| HP1 alpha | L929 | 2 | no barrier | 0.11 ± 0.03 | 0.03 ± 0.06 | 18 (100) |
| HP1 alpha | MTF | 2 | no barrier | 0.10 ± 0.03 | 0.05 ± 0.07 | 23 (100) |
| HP1 alpha | NIH3T3 | 2 | no barrier | 0.10 ± 0.05 | 0.05 ± 0.06 | 10 (100) |
| HP1 alpha | J1 NSC | 2 | no barrier | 0.10 ± 0.04 | 0.02 ± 0.12 | 15 (100) |
| HP1 beta | C2C12 | 2 | barrier | 0.53 ± 0.05 | 0.54 ± 0.14 | 13 (41.9) |
| HP1 beta | C2C12 |  | no barrier | 0.05 ± 0.02 | 0.04 ± 0.06 | 18 (58.1) |
| HP1 beta | L929 | 4 | barrier | 0.47 ± 0.08 | 0.40 ± 0.25 | 5 (23.8) |
| HP1 beta | L929 |  | no barrier | 0.05 ± 0.03 | 0.03 ± 0.10 | 16 (76.2) |
| HP1 beta | MTF | 2 | barrier | 0.46 ± 0.19 | 0.45 ± 0.50 | 3 (17.6) |
| HP1 beta | MTF |  | no barrier | 0.14 ± 0.03 | 0.08 ± 0.06 | 14 (82.4) |
| HP1 beta | NIH3T3 | 2 | barrier | 0.49 ± 0.07 | 0.35 ± 0.22 | 9 (39.1) |
| HP1 beta | NIH3T3 |  | no barrier | 0.11 ± 0.04 | 0.05 ± 0.09 | 14 (60.9) |
| HP1 beta | J1 NSC | 2 | barrier | 0.45 ± 0.11 | 0.53 ± 0.19 | 7 (33.3) |
| HP1 beta | J1 NSC |  | no barrier | 0.08 ± 0.03 | 0.06 ± 0.06 | 14 (66.7) |
| HP1 gamma | C2C12 | 2 | no barrier | 0.13 ± 0.05 | 0.03 ± 0.16 | 12 (100) |
| HP1 gamma | L929 | 3 | no barrier | 0.12 ± 0.03 | 0.03 ± 0.13 | 23 (100) |
| HP1 gamma | MTF | 2 | no barrier | 0.09 ± 0.04 | 0.02 ± 0.07 | 15 (100) |
| HP1 gamma | NIH3T3 | 2 | no barrier | 0.06 ± 0.03 | 0.03 ± 0.04 | 15 (100) |
| HP1 gamma | J1 NSC | 2 | no barrier | 0.09 ± 0.02 | 0.05 ± 0.12 | 8 (100) |
| Mbd1 | C2C12 | 2 | barrier | 0.49 ± 0.04 | 0.46 ± 0.30 | 30 (52.6) |
| Mbd1 | C2C12 |  | no barrier | 0.10 ± 0.03 | 0.03 ± 0.22 | 27 (47.4) |
| Mbd1 | L929 | 3 | barrier | 0.52 ± 0.02 | 0.44 ± 0.36 | 22 (57.9) |
| Mbd1 | L929 |  | no barrier | 0.12 ± 0.03 | 0.02 ± 0.23 | 16 (42.1) |
| Mbd1 | MTF | 2 | barrier | 0.53 ± 0.05 | 0.37 ± 0.30 | 7 (30.4) |
| Mbd1 | MTF |  | no barrier | 0.12 ± 0.03 | 0.12 ± 0.10 | 16 (69.6) |
| Mbd1 | NIH3T3 | 2 | barrier | 0.56 ± 0.02 | 0.62 ± 0.12 | 12 (40.0) |
| Mbd1 | NIH3T3 |  | no barrier | 0.11 ± 0.04 | 0.04 ± 0.13 | 18 (60.0) |
| Mbd1 | J1 NSC | 2 | barrier | 0.55 ± 0.03 | 0.42 ± 0.19 | 11 (52.4) |
| Mbd1 | J1 NSC |  | no barrier | 0.12 ± 0.04 | 0.03 ± 0.10 | 10 (47.6) |
| Mbd2 | C2C12 | 2 | barrier | 0.44 ± 0.07 | 0.40 ± 0.12 | 5 (33.3) |
| Mbd2 | C2C12 |  | no barrier | 0.07 ± 0.04 | 0.04 ± 0.09 | 10 (66.7) |
| Mbd2 | L929 | 3 | barrier | 0.52 ± 0.07 | 0.31 ± 0.31 | 9 (34.6) |
| Mbd2 | L929 |  | no barrier | 0.08 ± 0.03 | 0.05 ± 0.13 | 17 (65.4) |
| Mbd2 | MTF | 2 | barrier | 0.51 ± 0.03 | 0.40 ± 0.10 | 20 (60.6) |
| Mbd2 | MTF |  | no barrier | 0.16 ± 0.05 | 0.07 ± 0.08 | 13 (39.4) |
| Mbd2 | NIH3T3 | 3 | barrier | 0.52 ± 0.06 | 0.38 ± 0.15 | 11 (35.5) |
| Mbd2 | NIH3T3 |  | no barrier | 0.10 ± 0.03 | 0.00 ± 0.06 | 20 (64.5) |
| Mbd2 | J1 NSC | 2 | barrier | 0.49 ± 0.09 | 0.47 ± 0.16 | 6 (25.0) |
| Mbd2 | J1 NSC |  | no barrier | 0.09 ± 0.03 | 0.01 ± 0.10 | 18 (75.0) |
| Mbd4 | C2C12 | 3 | no barrier | 0.09 ± 0.04 | 0.02 ± 0.12 | 18 (100) |
| Mbd4 | L929 |  | no barrier | 0.11 ± 0.06 | 0.05 ± 0.09 | 7 (100) |
| Mbd4 | MTF |  | no barrier | 0.06 ± 0.06 | 0.03 ± 0.09 | 7 (100) |
| Mbd4 | NIH3T3 |  | no barrier | 0.10 ± 0.03 | 0.01 ± 0.05 | 28 (100) |
| Mbd4 | J1 NSC |  | no barrier | 0.10 ± 0.05 | 0.01 ± 0.08 | 12 (100) |
| MeCP2 | C2C12 | 3 | barrier | 0.51 ± 0.02 | 0.37 ± 0.17 | 19 (40.4) |
| MeCP2 | C2C12 |  | no barrier | 0.10 ± 0.03 | 0.01 ± 0.10 | 28 (59.6) |
| MeCP2 | L929 | 3 | barrier | 0.55 ± 0.01 | 0.55 ± 0.07 | 54 (71.1) |
| MeCP2 | L929 |  | no barrier | 0.06 ± 0.03 | 0.02 ± 0.04 | 22 (28.9) |
| MeCP2 | MTF | 3 | barrier | 0.53 ± 0.04 | 0.47 ± 0.12 | 14 (38.9) |
| MeCP2 | MTF |  | no barrier | 0.11 ± 0.02 | 0.00 ± 0.03 | 22 (61.1) |
| MeCP2 | NIH3T3 | 3 | barrier | 0.54 ± 0.02 | 0.51 ± 0.15 | 23 (76.7) |
| MeCP2 | NIH3T3 |  | no barrier | 0.10 ± 0.08 | 0.02 ± 0.13 | 7 (23.3) |
| MeCP2 | J1 NSC | 3 | barrier | 0.50 ± 0.07 | 0.42 ± 0.18 | 6 (24.0) |
| MeCP2 | J1 NSC |  | no barrier | 0.09 ± 0.03 | 0.01 ± 0.04 | 19 (76.0) |

n: number of heterochromatin compartments and its percentage (in parenthesis) to the total explored

**Supplementary Table S6.** Dip values (in arbitrary units) from Figure 2 and S3, provided as mean ± 95% confidence interval.

| POI | cell line | replicates | condition | dip (individual) | dip (average) | n (%) |
| --- | --- | --- | --- | --- | --- | --- |
| GFP | hTERT-RPE1 | 2 | no barrier | 0.08 ± 0.03 | 0.08 ± 0.06 | 14 (100) |
| GFP | NDF | 2 | no barrier | 0.06 ± 0.02 | 0.06 ± 0.11 | 27 (100) |
| MaSat | hTERT-RPE1 | 4 | no barrier | 0.11 ± 0.06 | 0.03 ± 0.09 | 6 (100) |
| MaSat | NDF | 2 | no barrier | 0.06 ± 0.03 | 0.06 ± 0.06 | 10 (100) |
| H1.0 | hTERT-RPE1 | 2 | barrier | 0.52 ± 0.09 | 0.42 ± 0.16 | 6 (14.0) |
| H1.0 | hTERT-RPE1 |  | no barrier | 0.08 ± 0.03 | 0.02 ± 0.11 | 37 (86.0) |
| H1.0 | NDF | 2 | barrier | 0.47 ± 0.14 | 0.34 ± 0.23 | 6 (26.1) |
| H1.0 | NDF |  | no barrier | 0.06 ± 0.01 | 0.00 ± 0.04 | 17 (73.9) |
| H1.4 | hTERT-RPE1 | 2 | barrier | 0.47 ± 0.12 | 0.31 ± 0.09 | 3 (17.6) |
| H1.4 | hTERT-RPE1 |  | no barrier | 0.05 ± 0.03 | 0.09 ± 0.05 | 14 (82.4) |
| H1.4 | NDF | 2 | barrier | $0.48 \pm 0.13$ | $0.37 \pm 0.17$ | 6 (25.0) |
| H1.4 | NDF | | no barrier | $0.07 \pm 0.01$ | $0.01 \pm 0.02$ | 16 (75.0) |
| HP1alpha | hTERT-RPE1 | 6 | barrier | $0.51 \pm 0.02$ | $0.49 \pm 0.16$ | 34 (39.1) |
| HP1alpha | hTERT-RPE1 | | no barrier | $0.07 \pm 0.01$ | $0.03 \pm 0.04$ | 53 (60.9) |
| HP1alpha | NDF | 4 | barrier | $0.46 \pm 0.15$ | $0.38 \pm 0.25$ | 5 (15.2) |
| HP1alpha | NDF | | no barrier | $0.05 \pm 0.02$ | $0.01 \pm 0.02$ | 28 (84.8) |
| HP1beta | hTERT-RPE1 | 3 | no barrier | $0.09 \pm 0.05$ | $0.01 \pm 0.07$ | 13 (100) |
| HP1beta | NDF | 2 | no barrier | $0.06 \pm 0.05$ | $0.08 \pm 0.08$ | 8 (100) |
| HP1gamma | hTERT-RPE1 | 3 | no barrier | $0.09 \pm 0.03$ | $0.03 \pm 0.08$ | 15 (100) |
| HP1gamma | NDF | 2 | no barrier | $0.04 \pm 0.02$ | $0.00 \pm 0.02$ | 20 (100) |
| Mbd1 | hTERT-RPE1 | 4 | barrier | $0.50 \pm 0.06$ | $0.40 \pm 0.25$ | 11 (42.3) |
| Mbd1 | hTERT-RPE1 | | no barrier | $0.13 \pm 0.04$ | $0.02 \pm 0.11$ | 15 (57.7) |
| Mbd1 | NDF | 2 | barrier | $0.50 \pm 0.10$ | $0.30 \pm 0.02$ | 8 (33.3) |
| Mbd1 | NDF | | no barrier | $0.05 \pm 0.03$ | $0.02 \pm 0.06$ | 16 (66.6) |
| Mbd2 | hTERT-RPE1 | 2 | no barrier | $0.05 \pm 0.03$ | $0.00 \pm 0.01$ | 18 (100) |
| Mbd2 | NDF | 4 | barrier | $0.52 \pm 0.21$ | $0.30 \pm 0.02$ | 3 (11.1) |
| Mbd2 | NDF | | no barrier | $0.03 \pm 0.01$ | $0.02 \pm 0.02$ | 24 (88.9) |
| Mbd4 | hTERT-RPE1 | 2 | no barrier | $0.11 \pm 0.04$ | $0.04 \pm 0.07$ | 11 (100) |
| Mbd4 | NDF | | no barrier | $0.06 \pm 0.02$ | $0.00 \pm 0.02$ | 15 (100) |
| MeCP2 | hTERT-RPE1 | 4 | barrier | $0.45 \pm 0.06$ | $0.38 \pm 0.26$ | 5 (13.5) |
| MeCP2 | hTERT-RPE1 | | no barrier | $0.07 \pm 0.02$ | $0.00 \pm 0.04$ | 32 (86.5) |
| MeCP2 | NDF | 7 | barrier | $0.43 \pm 0.12$ | $0.25 \pm 0.39$ | 8 (18.6) |
| MeCP2 | NDF | | no barrier | $0.05 \pm 0.02$ | $0.01 \pm 0.02$ | 35 (11.4) |

n: number of heterochromatin compartments and its percentage (in parenthesis) to the total explored

**Supplementary Table S7.** Enrichment and heterochromatin size values (mean ± 95% CI) for the mouse and human screenings (Figures 1 and 2) and its correlation with the dip values (Figure S6).

| protein | cell line | enrichment | correlation | heterochromatin size | correlation |
| --- | --- | --- | --- | --- | --- |
| GFP | C2C12 | $1.0 \pm 0.1$ | -0.07 | $1.977 \pm 0.394$ | 0.29 |
| GFP | L929 | $1.5 \pm 0.3$ | 0.02 | $1.881 \pm 0.250$ | -0.22 |
| GFP | MTF | $1.0 \pm 0.1$ | 0.41 | $1.512 \pm 0.374$ | 0.08 |
| GFP | NIH-3T3 | $1.0 \pm 0.1$ | -0.46 | $1.627 \pm 0.446$ | 0.46 |
| GFP | NSC | $1.0 \pm 0.1$ | -0.47 | $1.912 \pm 0.766$ | 0.35 |
| GFP | hTERT-RPE1 | 1.2 ± 0.1 | 0.07 | 0.677 ± 0.140 | 0.36 |
| GFP | NDF | 1.2 ± 0.1 | -0.33 | 2.932 ± 0.377 | 0.21 |
| MaSat | C2C12 | 3.9 ± 0.5 | 0.03 | 1.635 ± 0.275 | 0.42 |
| MaSat | L929 | 3.2 ± 0.4 | -0.12 | 1.978 ± 0.194 | 0.16 |
| MaSat | MTF | 3.0 ± 0.9 | -0.42 | 1.322 ± 0.204 | 0.45 |
| MaSat | NIH-3T3 | 2.7 ± 0.3 | -0.07 | 1.593 ± 0.329 | 0.54 |
| MaSat | NSC | 2.9 ± 0.6 | 0.31 | 1.501 ± 0.467 | 0.24 |
| MaSat | hTERT-RPE1 | 1.1 ± 0.3 | 0.22 | 1.061 ± 0.470 | 0.21 |
| MaSat | NDF | 1.3 ± 0.3 | 0.34 | 0.698 ± 0.293 | -0.04 |
| H1.0 | C2C12 | 2.6 ± 0.3 | 0.07 | 1.725 ± 0.283 | 0.69 |
| H1.0 | L929 | 2.9 ± 0.3 | -0.09 | 1.813 ± 0.193 | 0.34 |
| H1.0 | MTF | 2.6 ± 0.4 | 0.00 | 1.483 ± 0.174 | 0.29 |
| H1.0 | NIH-3T3 | 2.3 ± 0.1 | 0.27 | 1.696 ± 0.243 | 0.51 |
| H1.0 | NSC | 2.9 ± 0.3 | -0.05 | 1.682 ± 0.216 | -0.02 |
| H1.0 | hTERT-RPE1 | 3.3 ± 0.3 | 0.23 | 1.290 ± 0.227 | -0.10 |
| H1.0 | NDF | 3.4 ± 0.7 | 0.25 | 1.412 ± 0.171 | -0.08 |
| H1.4 | C2C12 | 2.7 ± 0.2 | 0.05 | 1.476 ± 0.172 | 0.51 |
| H1.4 | L929 | 3.1 ± 0.7 | 0.05 | 1.671 ± 0.337 | 0.20 |
| H1.4 | MTF | 2.6 ± 0.3 | 0.05 | 1.488 ± 0.280 | 0.54 |
| H1.4 | NIH-3T3 | 2.4 ± 0.1 | 0.28 | 2.055 ± 0.275 | 0.50 |
| H1.4 | NSC | 3.3 ± 0.6 | 0.30 | 1.460 ± 0.262 | 0.51 |
| H1.4 | hTERT-RPE1 | 3.1 ± 0.5 | 0.46 | 1.172 ± 0.293 | -0.30 |
| H1.4 | NDF | 4.1 ± 0.8 | 0.33 | 1.055 ± 0.111 | 0.11 |
| HP1alpha | C2C12 | 2.9 ± 0.2 | -0.07 | 1.677 ± 0.303 | -0.35 |
| HP1alpha | L929 | 3.0 ± 0.8 | 0.17 | 1.544 ± 0.294 | 0.10 |
| HP1alpha | MTF | 3.7 ± 0.4 | 0.10 | 1.256 ± 0.225 | 0.36 |
| HP1alpha | NIH-3T3 | 3.9 ± 0.8 | -0.26 | 1.475 ± 0.350 | 0.82 |
| HP1alpha | NSC | 3.7 ± 0.5 | -0.33 | 1.093 ± 0.168 | -0.21 |
| HP1alpha | hTERT-RPE1 | 4.5 ± 0.6 | 0.30 | 0.895 ± 0.165 | -0.03 |
| HP1alpha | NDF | 8.2 ± 2.0 | 0.35 | 0.972 ± 0.231 | 0.11 |
| HP1beta | C2C12 | 5.3 ± 0.8 | 0.31 | 1.470 ± 0.245 | 0.04 |
| HP1beta | L929 | 6.1 ± 1.3 | 0.34 | 2.561 ± 0.454 | -0.25 |
| HP1beta | MTF | 5.8 ± 1.2 | -0.08 | 1.619 ± 0.389 | 0.51 |
| HP1beta | NIH-3T3 | 7.1 ± 0.7 | -0.20 | 1.490 ± 0.272 | 0.63 |
| HP1beta | NSC | 5.1 ± 0.7 | 0.26 | 1.645 ± 0.307 | 0.31 |
| HP1beta | hTERT-RPE1 | 5.4 ± 1.0 | 0.35 | 2.326 ± 1.384 | 0.25 |
| HP1beta | NDF | 5.0 ± 1.2 | -0.37 | 0.703 ± 0.199 | -0.19 |
| HP1gamma | C2C12 | 2.7 ± 0.3 | -0.35 | 2.074 ± 0.545 | 0.75 |
| HP1gamma | L929 | 3.4 ± 0.4 | 0.14 | 1.204 ± 0.160 | -0.07 |
| HP1gamma | MTF | 3.1 ± 0.4 | -0.15 | 1.137 ± 0.258 | 0.33 |
| HP1gamma | NIH-3T3 | 2.8 ± 0.3 | 0.53 | 1.299 ± 0.346 | 0.45 |
| HP1gamma | NSC | 4.0 ± 0.7 | -0.20 | 1.548 ± 0.767 | 0.43 |
| HP1gamma | hTERT-RPE1 | 3.6 ± 0.7 | -0.29 | 0.815 ± 0.205 | -0.05 |
| HP1gamma | NDF | 5.0 ± 1.4 | -0.11 | 0.809 ± 0.176 | 0.07 |
| Mbd1 | C2C12 | 6.1 ± 0.5 | -0.04 | 1.272 ± 0.124 | 0.38 |
| Mbd1 | L929 | 6.0 ± 0.6 | 0.05 | 1.137 ± 0.126 | 0.28 |
| Mbd1 | MTF | 5.6 ± 0.8 | 0.24 | 0.962 ± 0.165 | 0.43 |
| Mbd1 | NIH-3T3 | 6.2 ± 0.7 | -0.01 | 1.512 ± 0.194 | 0.67 |
| Mbd1 | NSC | 6.3 ± 0.8 | 0.40 | 1.518 ± 0.377 | 0.45 |
| Mbd1 | hTERT-RPE1 | 4.0 ± 0.7 | 0.29 | 1.203 ± 0.313 | 0.21 |
| Mbd1 | NDF | 5.5 ± 2.0 | 0.89 | 1.074 ± 0.265 | -0.38 |
| Mbd2 | C2C12 | 10.3 ± 2.2 | 0.77 | 1.779 ± 0.392 | 0.28 |
| Mbd2 | L929 | 9.5 ± 2.0 | 0.68 | 1.709 ± 0.388 | 0.60 |
| Mbd2 | MTF | 6.8 ± 0.6 | 0.04 | 2.110 ± 0.333 | 0.39 |
| Mbd2 | NIH-3T3 | 8.5 ± 1.2 | -0.09 | 1.713 ± 0.270 | 0.47 |
| Mbd2 | NSC | 11.0 ± 2.5 | -0.08 | 1.487 ± 0.385 | -0.22 |
| Mbd2 | hTERT-RPE1 | 3.7 ± 0.8 | 0.25 | 0.924 ± 0.199 | -0.05 |
| Mbd2 | NDF | 5.9 ± 1.5 | 0.00 | 1.400 ± 0.421 | 0.19 |
| Mbd4 | C2C12 | 5.0 ± 0.6 | -0.21 | 1.209 ± 0.230 | -0.28 |
| Mbd4 | L929 | 10.5 ± 5.9 | -0.06 | 1.029 ± 0.382 | -0.47 |
| Mbd4 | MTF | 3.8 ± 1.0 | 0.03 | 1.274 ± 0.385 | 0.33 |
| Mbd4 | NIH-3T3 | 3.9 ± 0.5 | 0.28 | 1.422 ± 0.324 | 0.34 |
| Mbd4 | NSC | 9.6 ± 3.8 | 0.13 | 1.303 ± 0.361 | -0.27 |
| Mbd4 | hTERT-RPE1 | 4.1 ± 0.8 | -0.62 | 0.755 ± 0.207 | 0.34 |
| Mbd4 | NDF | 3.2 ± 0.4 | 0.42 | 1.724 ± 0.498 | 0.62 |
| MeCP2 | C2C12 | 7.6 ± 0.6 | 0.33 | 1.725 ± 0.243 | 0.32 |
| MeCP2 | L929 | 7.3 ± 0.4 | 0.23 | 1.542 ± 0.137 | 0.25 |
| MeCP2 | MTF | 7.5 ± 0.9 | 0.58 | 0.988 ± 0.149 | 0.67 |
| MeCP2 | NIH-3T3 | 10.2 ± 1.1 | 0.16 | 1.636 ± 0.340 | 0.33 |
| MeCP2 | NSC | 9.4 ± 1.7 | -0.05 | 1.354 ± 0.264 | -0.31 |
| MeCP2 | hTERT-RPE1 | 2.0 ± 0.5 | 0.54 | 1.198 ± 0.353 | 0.16 |
| MeCP2 | NDF | 2.6 ± 0.5 | 0.16 | 2.680 ± 0.435 | -0.10 |

**Supplementary Table S8.** Dip values (in arbitrary units) from Figure 3, provided as mean ± 95% confidence interval.

| POI | cell line | replicates | condition | dip (individual) | dip (average) | n (%) |
| --- | --- | --- | --- | --- | --- | --- |
| Mbd1 | MEFp | 2 | barrier | 0.40 ± 0.04 | 0.33 ± 0.07 | 22 (66.7) |
| Mbd1 | MEFp |  | no barrier | 0.17 ± 0.04 | 0.13 ± 0.07 | 11 (33.3) |
| Mbd1 | MEFpm | 2 | barrier | 0.41 ± 0.05 | 0.34 ± 0.13 | 13 (44.8) |
| Mbd1 | MEFpm |  | no barrier | 0.13 ± 0.03 | 0.10 ± 0.03 | 16 (55.2) |
| Mbd2 | MEFp | 2 | barrier | 0.38 ± 0.05 | 0.31 ± 0.08 | 12 (44.4) |
| Mbd2 | MEFp |  | no barrier | 0.15 ± 0.03 | 0.12 ± 0.05 | 15 (55.6) |
| Mbd2 | MEFpm | 2 | barrier | 0.37 ± 0.08 | 0.23 ± 0.12 | 6 (40.0) |
| Mbd2 | MEFpm |  | no barrier | 0.14 ± 0.05 | 0.08 ± 0.04 | 9 (60.0) |
| MeCP2 | MEFp | 2 | barrier | 0.37 ± 0.03 | 0.34 ± 0.05 | 38 (62.3) |
| MeCP2 | MEFp |  | no barrier | 0.13 ± 0.04 | 0.12 ± 0.02 | 23 (41.7) |
| MeCP2 | MEFpm | 2 | barrier | 0.31 ± 0.06 | 0.26 ± 0.06 | 18 (31.0) |
| MeCP2 | MEFpm |  | no barrier | 0.11 ± 0.02 | 0.07 ± 0.02 | 30 (69.0) |
| HP1beta | MEF w8 | 2 | barrier | 0.40 ± 0.06 | 0.34 ± 0.09 | 13 (54.2) |
| HP1beta | MEF w8 |  | no barrier | 0.11 ± 0.04 | 0.07 ± 0.08 | 11 (45.8) |
| HP1beta | MEF D5 | 2 | barrier | 0.29 ± 0.02 | 0.25 ± 0.07 | 4 (50.0) |
| HP1beta | MEF D5 |  | no barrier | 0.09 ± 0.06 | 0.06 ± 0.04 | 4 (50.0) |

n: number of heterochromatin compartments and its percentage (in parenthesis) to the total explored

**Supplementary Table S9.** High-throughput microscopy analysis statistic from Figure 4. The intensity values are given as mean ± 95% confidence interval. The Hedge G is also having the 95% confidence interval.

| Cell line | replicate | n cells | staining | norm. int. (a.u.) | versus | welch p-value | Hedges G |
| --- | --- | --- | --- | --- | --- | --- | --- |
| C2C12 | R1 | 2998 | 5mC | 2.329 ± 0.009 | L929 | 3.7x10<sup>-5</sup> | -2.584 ± 0.826 |
| C2C12 | R2 | 2459 | 5mC | 2.456 ± 0.011 | MTF | 7.6x10<sup>-7</sup> | -3.673 ± 1.041 |
| C2C12 | R3 | 950 | 5mC | 1.544 ± 0.016 | NIH-3T3 | 9.1x10<sup>-4</sup> | -2.011 ± 0.690 |
| C2C12 | R4 | 3446 | 5mC | 2.357 ± 0.022 | NSC | 1.1x10<sup>-3</sup> | -4.985 ± 1.328 |
| C2C12 | R5 | 1780 | 5mC | 3.093 ± 0.031 |  |  |  |
| C2C12 | R6 | 1039 | 5mC | 2.287 ± 0.018 |  |  |  |
| C2C12 | R7 | 257 | 5mC | 2.491 ± 0.033 |  |  |  |
| C2C12 | R8 | 12127 | 5mC | 2.207 ± 0.013 |  |  |  |
| C2C12 | R9 | 6932 | 5mC | 2.796 ± 0.018 |  |  |  |
| C2C12 | R10 | 7499 | 5mC | 2.063 ± 0.014 |  |  |  |
| **C2C12** | **all** |  | **5mC** | **2.362 ± 0.296** |  |  |  |
| L929 | R1 | 4225 | 5mC | 3.721 ± 0.036 | C2C12 | 3.7x10<sup>-5</sup> | 2.584 ± 0.826 |
| L929 | R2 | 4220 | 5mC | 3.979 ± 0.034 | MTF | 0.35 | -0.469 ± 1.168 |
| L929 | R3 | 15350 | 5mC | 3.460 ± 0.019 | NIH-3T3 | 0.70 | 0.168 ± 1.280 |
| L929 | R4 | 307 | 5mC | 3.105 ± 0.052 | NSC | 0.013 | -2.439 ± 0.606 |
| L929 | R5 | 1923 | 5mC | 3.060 ± 0.047 |  |  |  |
| L929 | R6 | 259 | 5mC | 2.906 ± 0.042 |  |  |  |
| L929 | R7 | 1110 | 5mC | 3.679 ± 0.033 |  |  |  |
| L929 | R8 | 622 | 5mC | 3.867 ± 0.035 |  |  |  |
| **L929** | **all** |  | **5mC** | **3.472 ± 0.338** |  |  |  |
| MTF | R1 | 3021 | 5mC | 3.469 ± 0.035 | C2C12 | 7.6x10<sup>-7</sup> | 3.673 ± 1.041 |
| MTF | R2 | 13004 | 5mC | 3.739 ± 0.030 | L929 | 0.35 | 0.469 ± 1.168 |
| MTF | R3 | 2053 | 5mC | 3.410 ± 0.042 | NIH-3T3 | 0.26 | 0.592 ± 1.680 |
| MTF | R4 | 1547 | 5mC | 3.198 ± 0.023 | NSC | 0.037 | -3.378 ± 3.294 |
| MTF | R5 | 2370 | 5mC | 3.678 ± 0.020 |  |  |  |
| MTF | R6 | 108 | 5mC | 3.580 ± 0.146 |  |  |  |
| MTF | R7 | 1318 | 5mC | 3.5698 ± 0.029 |  |  |  |
| MTF | R8 | 599 | 5mC | 3.768 ± 0.037 |  |  |  |
| **MTF** | **all** |  | **5mC** | **3.568 ± 0.164** |  |  |  |
| NIH-3T3 | R1 | 10523 | 5mC | 3.078 ± 0.017 | C2C12 | 9.1x10<sup>-4</sup> | 2.011 ± 0.690 |
| NIH-3T3 | R2 | 7072 | 5mC | 2.700 ± 0.056 | L929 | 0.70 | -0.168 ± 1.280 |
| NIH-3T3 | R3 | 4033 | 5mC | 3.916 ± 0.038 | MTF | 0.26 | -0.592 ± 1.680 |
| NIH-3T3 | R4 | 890 | 5mC | 2.983 ± 0.051 | NSC | 6.9x10<sup>-3</sup> | -2.024 ± 2.704 |
| NIH-3T3 | R5 | 666 | 5mC | 3.754 ± 0.111 |  |  |  |
| NIH-3T3 | R6 | 92 | 5mC | 4.283 ± 0.189 |  |  |  |
| NIH-3T3 | R7 | 529 | 5mC | 3.370 ± 0.052 |  |  |  |
| NIH-3T3 | R8 | 251 | 5mC | 2.930 ± 0.097 |  |  |  |
| **NIH-3T3** | **all** |  | **5mC** | **3.377 ± 0.464** |  |  |  |
| NSC | R1 | 18330 | 5mC | 4.930 ± 0.021 | C2C12 | 1.1x10<sup>-3</sup> | 4.985 ± 1.328 |
| NSC | R2 | 17915 | 5mC | 4.323 ± 0.019 | L929 | 0.013 | 2.439 ± 0.606 |
| NSC | R3 | 2630 | 5mC | 4.312 ± 0.059 | MTF | 0.037 | 3.378 ± 3.294 |
| **NSC** | **all** |  | **5mC** | **4.522 ± 0.879** | NIH-3T3 | 6.9x10<sup>-3</sup> | 2.024 ± 2.704 |
| RPE-1 | R1 | 3046 | 5mC | 5.317 ± 0.044 | NDF | 0.704 | 0.0257 ± 1.086 |
| RPE-1 | R2 | 3696 | 5mC | 4.052 ± 0.032 |  |  |  |
| RPE-1 | R3 | 6005 | 5mC | 4.493 ± 0.031 |  |  |  |
| RPE-1 | R4 | 5606 | 5mC | 4.855 ± 0.027 |  |  |  |
| RPE-1 | R5 | 768 | 5mC | 5.034 ± 0.078 |  |  |  |
| RPE-1 | R6 | 672 | 5mC | 4.208 ± 0.074 |  |  |  |
| RPE-1 | R7 | 599 | 5mC | 4.181 ± 0.077 |  |  |  |
| RPE-1 | R8 | 875 | 5mC | 4.623 ± 0.087 |  |  |  |
| RPE-1 | R9 | 873 | 5mC | 5.164 ± 0.073 |  |  |  |
| RPE-1 | R10 | 826 | 5mC | 5.131 ± 0.082 |  |  |  |
| **RPE-1** | **all** |  | **5mC** | **4.706 ± 0.328** |  |  |  |
| NDF | R1 | 402 | 5mC | 4.388 ± 0.148 | RPE | 0.704 | -0.0257 ± 1.086 |
| NDF | R2 | 456 | 5mC | 4.028 ± 0.112 |  |  |  |
| NDF | R3 | 2262 | 5mC | 4.896 ± 0.042 |  |  |  |
| NDF | R4 | 1872 | 5mC | 5.064 ± 0.049 |  |  |  |
| **NDF** | **all** |  | **5mC** | **4.594 ± 0.755** |  |  |  |
| C2C12 | R1 | 3639 | H3K9me3 | 0.440 ± 0.004 | L929 | 1.6x10<sup>-4</sup> |  |
| C2C12 | R2 | 3195 | H3K9me3 | 0.391 ± 0.004 | MTF | 1.0x10<sup>-7</sup> |  |
| C2C12 | R3 | 1886 | H3K9me3 | 0.466 ± 0.006 | NIH-3T3 | 1.1x10<sup>-3</sup> |  |
| C2C12 | R4 | 613 | H3K9me3 | 0.511 ± 0.011 | NSC | 1.0x10<sup>-3</sup> |  |
| C2C12 | R5 | 414 | H3K9me3 | 0.517 ± 0.016 |  |  |  |
| C2C12 | R6 | 1015 | H3K9me3 | 0.461 ± 0.007 |  |  |  |
| C2C12 | R7 | 522 | H3K9me3 | 0.540 ± 0.014 |  |  |  |
| C2C12 | R8 | 462 | H3K9me3 | 0.360 ± 0.010 |  |  |  |
| C2C12 | R9 | 1883 | H3K9me3 | 0.453 ± 0.06 |  |  |  |
| C2C12 | R10 | 1003 | H3K9me3 | 0.465 ± 0.008 |  |  |  |
| C2C12 | R11 | 1313 | H3K9me3 | 0.451 ± 0.007 |  |  |  |
| **C2C12** | **all** |  | **H3K9me3** | **0.460 ± 0.035** |  |  |  |
| L929 | R1 | 1285 | H3K9me3 | 1.119 ± 0.020 | C2C12 | 1.6x10<sup>-4</sup> |  |
| L929 | R2 | 2009 | H3K9me3 | 1.038 ± 0.014 | MTF | 1.9x10<sup>-3</sup> |  |
| L929 | R3 | 2151 | H3K9me3 | 0.785 ± 0.010 | NIH-3T3 | 0.114 |  |
| L929 | R4 | 325 | H3K9me3 | 1.019 ± 0.041 | NSC | 3.0x10<sup>-5</sup> |  |
| L929 | R5 | 599 | H3K9me3 | 1.083 ± 0.024 |  |  |  |
| L929 | R6 | 2081 | H3K9me3 | 0.598 ± 0.008 |  |  |  |
| L929 | R7 | 83 | H3K9me3 | 0.596 ± 0.032 |  |  |  |
| L929 | R8 | 121 | H3K9me3 | 1.104 ± 0.059 |  |  |  |
| L929 | R9 | 103 | H3K9me3 | 0.850 ± 0.055 |  |  |  |
| **L929** | **all** |  | **H3K9me3** | **0.910 ± 0.162** |  |  |  |
| MTF | R1 | 2094 | H3K9me3 | 1.107 ± 0.016 | C2C12 | 1.0x10<sup>-7</sup> |  |
| MTF | R2 | 1194 | H3K9me3 | 1.163 ± 0.019 | L929 | 1.9x10<sup>-3</sup> |  |
| MTF | R3 | 694 | H3K9me3 | 1.230 ± 0.024 | NIH-3T3 | 0.165 |  |
| MTF | R4 | 913 | H3K9me3 | 1.160 ± 0.019 | NSC | 1.7x10<sup>-4</sup> |  |
| MTF | R5 | 1921 | H3K9me3 | 1.165 ± 0.014 |  |  |  |
| MTF | R6 | 1818 | H3K9me3 | 1.574 ± 0.020 |  |  |  |
| MTF | R7 | 6602 | H3K9me3 | 1.022 ± 0.007 |  |  |  |
| MTF | R8 | 3206 | H3K9me3 | 1.541 ± 0.015 |  |  |  |
| MTF | R9 | 782 | H3K9me3 | 1.379 ± 0.023 |  |  |  |
| MTF | R10 | 681 | H3K9me3 | 1.153 ± 0.023 |  |  |  |
| **MTF** | **all** |  | **H3K9me3** | **1.249 ± 0.133** |  |  |  |
| NIH-3T3 | R1 | 1453 | H3K9me3 | 0.944 ± 0.012 | C2C12 | 1.1x10<sup>-3</sup> |  |
| NIH-3T3 | R2 | 564 | H3K9me3 | 0.903 ± 0.033 | L929 | 0.114 |  |
| NIH-3T3 | R3 | 85 | H3K9me3 | 1.178 ± 0.065 | MTF | 0.165 |  |
| NIH-3T3 | R4 | 300 | H3K9me3 | 1.350 ± 0.045 | NSC | 3.4x10<sup>-5</sup> |  |
| NIH-3T3 | R5 | 98 | H3K9me3 | 1.110 ± 0.043 |  |  |  |
| **NIH-3T3** | **all** |  | **H3K9me3** | **1.097 ± 0.225** |  |  |  |
| NSC | R1 | 294 | H3K9me3 | 3.109 ± 0.190 | C2C12 | 1.0x10<sup>-3</sup> |  |
| NSC | R2 | 473 | H3K9me3 | 3.272 ± 0.096 | L929 | 3.0x10<sup>-5</sup> |  |
| NSC | R3 | 106 | H3K9me3 | 2.933 ± 0.207 | MTF | 1.7x10<sup>-4</sup> |  |
| **NSC** | **all** |  | **H3K9me3** | **3.105 ± 0.420** | NIH-3T3 | 3.4x10<sup>-5</sup> |  |
| RPE-1 | R1 | 4744 | H3K9me3 | 4.514 ± 0.032 | NDF | 0.036 | -4.8102 ± 1.042 |
| RPE-1 | R2 | 3696 | H3K9me3 | 4.052 ± 0.032 |  |  |  |
| RPE-1 | R3 | 6854 | H3K9me3 | 3.890 ± 0.023 |  |  |  |
| RPE-1 | R4 | 6926 | H3K9me3 | 4.009 ± 0.024 |  |  |  |
| **RPE-1** | **all** |  | **H3K9me3** | **4.116 ± 0.436** |  |  |  |
| NDF | R1 | 2179 | H3K9me3 | 6.114 ± 0.083 | RPE1 | 0.036 | -4.8102 ± 1.042 |
| NDF | R2 | 2436 | H3K9me3 | 5.632 ± 0.069 |  |  |  |
| **NDF** | **all** |  | **H3K9me3** | **5.873 ± 3.061** |  |  |  |

**Supplementary Table S10.** Statistics for the Pearson correlation coefficient in Figure 4.

| cell line | replicate | n cells 5mC | PCC 5mC | n cells H3K9me3 | PCC H3K9me3 | p-value PCC 5mc<br>vs H3K9me3 |
| --- | --- | --- | --- | --- | --- | --- |
| C2C12 | R1 | 16 | $0.674 \pm 0.058$ | 8 | $0.659 \pm 0.066$ | |
| C2C12 | R2 | 17 | $0.684 \pm 0.056$ | 13 | $0.692 \pm 0.040$ | |
| C2C12 | R3 | 11 | $0.715 \pm 0.044$ | 19 | $0.706 \pm 0.025$ | |
| **C2C12** | **all** | **44** | **$0.691 \pm 0.053$** | **40** | **$0.686 \pm 0.060$** | **0.798 (n.s.)** |
| L929 | R1 | 7 | $0.724 \pm 0.089$ | 7 | $0.649 \pm 0.047$ | |
| L929 | R2 | 4 | $0.684 \pm 0.081$ | 8 | $0.652 \pm 0.034$ | |
| L929 | R3 | 6 | $0.715 \pm 0.068$ | 10 | $0.566 \pm 0.086$ | |
| **L929** | **all** | **17** | **$0.708 \pm 0.066$** | **25** | **$0.622 \pm 0.121$** | **0.055 (n.s.)** |
| MTF | R1 | 10 | $0.660 \pm 0.092$ | 10 | $0.688 \pm 0.070$ | |
| MTF | R2 | 7 | $0.597 \pm 0.107$ | 7 | $0.652 \pm 0.135$ | |
| MTF | R3 | 9 | $0.646 \pm 0.072$ | 9 | $0.566 \pm 0.065$ | |
| **MTF** | **all** | **26** | **$0.634 \pm 0.083$** | **20** | **$0.708 \pm 0.134$** | **0.114 (n.s.)** |
| NIH-3T3 | R1 | 9 | $0.611 \pm 0.071$ | 9 | $0.709 \pm 0.051$ | |
| NIH-3T3 | R2 | 3 | $0.576 \pm 0.271$ | 3 | $0.708 \pm 0.063$ | |
| NIH-3T3 | R3 | 9 | $0.729 \pm 0.039$ | 9 | $0.689 \pm 0.079$ | |
| NIH-3T3 | R4 | 6 | $0.612 \pm 0.139$ | 6 | $0.643 \pm 0.069$ | |
| **NIH-3T3** | **all** | **27** | **$0.632 \pm 0.106$** | **29** | **$0.687 \pm 0.049$** | **0.185 (n.s.)** |
| NSC | R1 | 5 | $0.608 \pm 0.140$ | 4 | $0.683 \pm 0.191$ | |
| NSC | R2 | 5 | $0.756 \pm 0.073$ | 3 | $0.679 \pm 0.246$ | |
| NSC | R3 | | | 3 | $0.686 \pm 0.231$ | |
| NSC | R4 | | | 4 | $0.679 \pm 0.101$ | |
| **NSC** | **all** | **10** | **$0.682 \pm 0.943$** | **14** | **$0.683 \pm 0.009$** | **0.999 (n.s.)** |
| hTERT-RPE1 | R1 | 13 | $0.227 \pm 0.098$ | 30 | $0.481 \pm 0.038$ | |
| hTERT-RPE1 | R2 | 22 | $0.293 \pm 0.082$ | 17 | $0.567 \pm 0.055$ | |
| hTERT-RPE1 | R3 | 27 | $0.174 \pm 0.088$ | 21 | $0.496 \pm 0.050$ | |
| hTERT-RPE1 | R4 | 30 | $0.097 \pm 0.075$ | | | |
| **hTERT-RPE1** | **all** | **79** | **$0.198 \pm 0.132$** | **68** | **$0.515 \pm 0.114$** | **0.002 (\*)** |
| NDF | R1 | 12 | $0.340 \pm 0.134$ | 13 | $0.474 \pm 0.152$ | |
| NDF | R2 | 9 | $0.222 \pm 0.100$ | 18 | $0.588 \pm 0.134$ | |
| NDF | R3 | | | 14 | $0.441 \pm 0.133$ | |
| **NDF** | **all** | **21** | **$0.281 \pm 0.751$** | **45** | **$0.501 \pm 0.191$** | **0.056 (n.s.)** |

## Notes

### Competing Interest Statement

The authors have declared no competing interest.

